# Triple-action inhibitory mechanism of allosteric TYK2-specific inhibitors

**DOI:** 10.1101/2023.10.09.561507

**Authors:** Jimin Wang, Victor S. Batista, Christopher G. Bunick

## Abstract

Deucravacitinib, 6-(cyclopropanecarbonylamido)-4-[2-methoxy-3-(1-methyl-1,2,4-triazol-3-yl)anilino]-N-(trideuteriomethyl)pyridazine-3-carboxamide, is a highly selective inhibitor of protein tyrosine kinase 2 (TYK2) that targets the Janus kinase (JAK)-signal transducer and activator of transcription (STAT) pathway. The structural basis for its selectivity and allosteric inhibition remains poorly understood. Here, we investigate the inhibition mechanism through analysis of available structures relevant to the STAT pathway, including crystal structures of the truncated TYK2 FERM-SH2 domain bound to the IFNα type I receptor (IFNαR1) and the truncated TYK2 JH2-JH1 domain. Our computational analysis provides a mechanistic hypothesis for the relatively rapid interferon-induced gene expression mediated by TYK2 relative to other cytokines. We find that deucravacitinib inhibits TYK2 kinase in three distinct states: the autoinhibited state and two activated states for autophosphorylation and phosphorylation of downstream protein substrates. Its binding to the TYK2 pseudokinase domain in the autoinhibited state restricts the essential dynamics of the TYK2 kinase domain required for kinase activity. Furthermore, it binds competitively with ATP in the pseudokinase domain, and also directly prevents formation of the active state of TYK2 through steric clashes.

## Introduction

Interferons (IFNs) are a class of cytokines produced and released by host cells in response to bacterial and viral infections. These small proteins activate a large number of *interferon-stimulated genes* (ISG) that are essential for a rapid immune response. IFNs, therefore, are essential for the organism immune response to an infection. Here, we report a structural analysis that provides fundamental insights on the mechanism of allosteric inhibition of protein tyrosine kinase 2 (TYK2) by the highly selective inhibitor deucravacitinib (DEU; marketed under brand name Sotyktu), i.e, 6-(cyclopropanecarbonylamido)-4-[2-methoxy-3-(1-methyl-1,2,4-triazol-3-yl)anilino]-N-(trideuteriomethyl)pyridazine-3-carboxamide.

The non-receptor protein tyrosine kinase 2 (TYK2) was first identified and cloned in 1990 (Firmbach-Kraft et al., 1990; Krolewski et al., 1990). Two years later, JAK1 and JAK2 were discovered and named after the double-faced Roman God Janus due to the tandem repeat in the primary sequence of one pseudokinase and one kinase domain [see (Philips et al., 2022) for a comprehensive account of the discovery of the JAK-STAT signaling pathway]. However, it took ∼30 years to uncover the first structure of the activated JAK1 homodimeric complex with non-physiological chimeric interferon-λ type I receptor (IFNλR1) domains (Caveney et al., 2023; Glassman et al., 2022), partially due to the nature of highly dynamical proteins with multiple functional states. Progress in structural elucidation has occurred using ADP to reduce the heterogeneity of partial phosphorylation of the enzymes and receptors. Using the chimeric receptor, which is fully capable of JAK1 kinase autophosphorylation, it was possible to simplify the activated complex of two receptors and two kinases to only two proteins of a receptor and a kinase.

Inhibitors like DEU selectively inhibit only TYK2 but not the 3 other JAK enzymes, suggesting that key structural differences exist among them. The hallmark of this inhibitor class is they bind JH2 (JAK homolog domain) but allosterically inhibit JH1 in mechanisms that are not fully understood. Here, we apply homology modeling procedures to investigate the heterodimeric TYK2/JAK1 complex in response to IFNα and IFNγ and the potential regulatory mechanism of specific inhibitors of TYK2 JH2. Our full-length model of the TYK2 structure is built from the crystal structures of the truncated TYK2 FERM-SH2 domain bound to the IFNα type I receptor (IFNαR1) as well as the truncated TYK2 JH2-JH1 domain (Lupardus et al., 2014; Wallweber et al., 2014).

All protein kinases have 11 conserved motifs for folding of a bilobal structure, including a β-barrel like N-terminal lobe, and a helical C-terminal lobe (Fig. 1) (Hanks et al., 1988). ATP and protein substrates bind to the cleft between the two lobes, exemplified by the IRTYK complex with a bi-substrate analog inhibitor (Protein Data Bank, PDB ID code 1gag) (Parang et al., 2001). Kinase domains have a conserved catalytic triad, including a Lys residue in the VAVK sequence motif that orients the ATP triphosphate moiety by forming a H bond with its α- phosphate group, an Asp residue in the HRD motif that removes a proton from the Tyr sidechain to make it a nucleophile, and another Asp residue in the DFG motif (together with an Asn) that binds the two divalent metal ions that orient the γ-phosphate. Importantly, pseudokinase domains are missing some of the 3 catalytic residues. Kinases also include a doublet phosphorylated tyrosine motif (pYpY) which serves as a docking site for downstream protein substrates (receptors and uSTATs). These kinase domains have other phosphorylation sites of Tyr and other post-translational modifications (PTMs) that are often specific to a given JAK kinase but not always conserved. The access of γ-phosphate to the nucleophile is controlled by an opening-closing, or an up-down motion of the GXGXΦG β-hairpin motif where Φ is either Phe or Tyr residue. The adenine base is recognized by backbone H bonds of a short β-stretch beginning with Glu, in the (M978)-EYVF motif in TYK2 JH1, located on the backside of the cleft. This β-stretch exhibits a direct contact with TYK2 JH2 in the autoinhibited state (Lupardus et al., 2014). Different sequences are found in 3 other JAKs, including (M956)-EFLP in JAK1 JH1, (M929)-EYLP in JAK2 JH1, and (M902)-EpYLP in JAK3 JH1, which involves a phosphorylated pY. These differences might suggest different stability of the autoinhibited states of different JAK enzymes.

**Figure 1.**
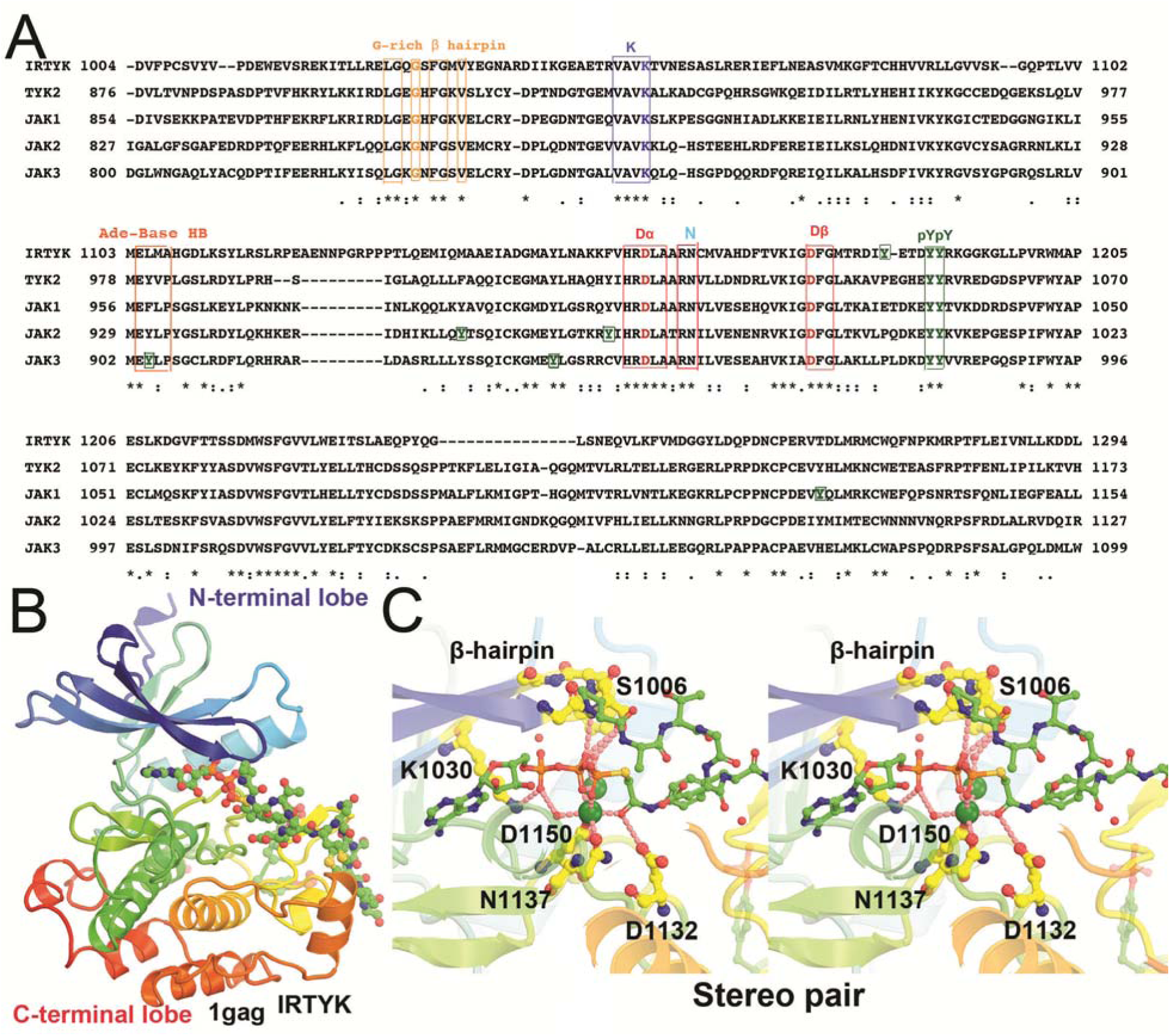
Kinase domain sequence alignment of IRTYK, TYK2, JAK1 (double-headed Janus kinase), JAK2, and JAK3 in context of the IRTYK complex with bi-substrate analog. (A) Sequence alignment. ATP-interacting motifs are boxed, including the β-hairpin of the GXGXΦG motif (a flip that controls the access of substrate to the γ-phosphate) where Φ is either F or Y, a catalytic triad K, D, and D in the VAVK, HRD, and DFG motifs, respectively, and the adenine base recognition loop of the MEΦXP motif of JAKs. The conserved substrate-docking doublet pY/pY site and other non-conserved single pY sites are also boxed in green. (B) Bilobal kinase domain in complex with the bi-substrate analog in rainbow colors from β-barrel like N-terminal lobe to helical C-terminal lobe. The bi-substrate analog in balls-and-sticks model. (C) Stereodiagram of the close-up view of the kinase active site in which two large green spheres represent two Mg^2+^ ions.

Protein kinases transfer a phosphate group from ATP to protein substrate. The catalytic subunits are defined by a smaller N-terminal lobe and a larger C-terminal lobe (Fig. 1). These two lobes are separated by a deep cleft for binding of Mg^2+^/ ATP and a groove for binding of substrate(s). ATP binding to the cleft between the two lobes precedes substrate binding because the phosphate group of ATP forms hydrogen bonds with hydroxyl groups (of Tyr, Ser or Thr) in the protein substrates before catalysis. A prominent example is the insulin receptor protein tyrosine kinase (IRTYK) bound to a bi-substrate analog inhibitor (Fig. 1) (Parang et al., 2001). All TYKs have the phosphotyrosine (pY) doublet as a substrate docking site after activation. The pY/pY doublet site is pY1054/pY1055 in TYK2 JH1, and pY1033/pY1034 in JAK1 JH1. A main difference between kinases and pseudokinases is that the latter lack one or more of the catalytic residues such as in the KDD catalytic triad and are thereby inactive (Fig 1). The human genome includes ∼538 protein kinases, which account for 2% of all human genes (Manning et al., 2002). Of these, 456 (85% of all kinases) have been specifically identified in cellular localization studies (Zhang et al., 2021). With so many kinase sequences known, JAK kinases were often dubbed as yet “Just Another Kinase” since they share a high degree of sequence and structural conservation with a bilobal fold (Hanks et al., 1988).

Studies of the JAK-STAT immune system signaling pathway elucidated how it contributes to immunity against bacterial and viral infections. This signaling pathway is also involved in autoimmune, inflammatory, and allergic diseases. Therefore, the JAK-STAT pathway is a promising target for drug discovery (Liu et al., 2021; Spinelli et al., 2021). In fact, drug discovery efforts over the past decade focused initially on targeting gain-of-function mutated JAK2. Here, we find that DEU exhibits an inhibitory effect in two distinct states of the TYK2 kinase domain. Its binding to the TYK2 pseudokinase domain in the autoinhibited state restricts the essential dynamics of the TYK2 kinase domain required for kinase activity. Furthermore, it binds competitively with ATP and directly prevents formation of the active state of TYK2.

When IFNs or other cytokines bind to their corresponding receptors, the intracellular domains of the receptors unfold, allowing JAKs to dimerize and activate via autophosphorylation. This has been reported for TYK2, JAK1, JAK2, and JAK3 in a variety of homo-and heterodimeric combinations, and even in higher oligomers (Glassman et al., 2022; Philips et al., 2022). JAKs have several domains, including the Four-point-one, Ezrin, Radixin, and Moesin (FERM) motif (JH7-JH4), SH2 (JH3), JH2, and JH1. They phosphorylate a selective set of 7 uSTAT proteins (STAT1-4, 5A, 5B, and 6), which share 6 domains of N-terminal (NTD), coiled-coil (CCD), DNA-binding (DBD), linker (LD), SH2, and transcriptional activation (TAD) domains (Becker et al., 1998; Chen et al., 1998). Their SH2 domain contains a phosphorylation tail segment for phosphorylation-dependent dimerization. Before Tyr phosphorylation, uSTAT forms an antiparallel dimer (La Sala et al., 2020; Mao et al., 2005; Neculai et al., 2005). After Tyr phosphorylation, pSTAT exists in a parallel dimer (Becker et al., 1998; Chen et al., 1998; Li et al., 2016), exposing nuclear localization signals for it to import into the nucleus in an importins-dependent manner (Ernst and Muller-Newen, 2019; Meyer and Vinkemeier, 2004). Some uSTAT1 import into and export out of the nucleus in a carrier-independent manner.

DEU allosterically inhibits autophosphorylation of 6 TYK2 Tyr residues: two in JH1, two in JH2, and two in the JH4-JH7 domains. Multiple pY sites on the activated TYK2 surface serve as recruitment stations for unphosphorylated STAT (uSTAT) protein substrates. Through a process of binding and transient release, uSTAT proteins migrate to the high affinity site that contains two adjacent pY1054/pY1055 doublet in JH1 to be phosphorylated to become pSTAT. After phosphorylation and dimerization of two pSTAT monomers, the dimeric pSTAT product is released and the doublet pY/pY site in JH1 is vacated for binding of the next uSTAT substrates. The process of initial binding of uSTAT monomers and then its moving onto the continuously vacated catalytic site in a conveyer belt-like architecture makes the TYK2 surface highly efficient for phosphorylating STATs and assembling pSTAT dimers.

Before tyrosine phosphorylation, vacant pY binding sites of the SH2 domain of uSTAT antiparallel dimers are attracted to the pY sites of the activated JAK complex for initial binding. Because the activated TYK2 has 6 pY sites (pY292, pY433, pY604, pY827, and pY1054/pY1055), more than other JAK enzymes (e.g., JAK1 has only 3, pY1033/pY1034, and pY1125), uSTAT substrates can be enriched selectively on TYK2 for rapid phosphorylation, as will be addressed below in this study. In addition, the pY sites of JAKs serve as recruitment stations for SH2 domains of negative competitive regulators such as SOCS (suppressor of cytokine signaling) family proteins. Other negative regulators include the PIAS (protein inhibitors of activated STAT) and PTP (protein tyrosine phosphatases) family proteins as well as ISG15 and USP18, which is a ubiquitin specific peptidase-18 – a deubiquitinating protease for removing ISG15 conjugate in a process known as delSGylation. Activated JAKs also phosphorylate tyrosine residues of cytokine receptors (Greenlund et al., 1994), which may play some role in selectivity for specific uSTAT proteins. Besides tyrosine phosphorylation, both JAKs and STATs have many other PTMs to integrate signals from other pathways and processes, including methylation, acetylation, sumoylation, threonine and serine phosphorylation. There are as many as 80 different types of PTMs for STAT3 (Diallo and Herrera, 2022). Therefore, STATs may play many important roles other than the canonical function of transcription (Stark and Darnell, 2012).

DEU is highly selective against TYK2 JH2 among a panel of 249 protein and lipid kinases and pseudokinases examined (Fig. 2) (Burke et al., 2019). Its median inhibitory concentration (IC_50_) against TYK2 JH2 is 0.2 *n*M. It has some minor effects on only two other molecules among this panel tested, having IC_50_ of less than 200 nM: one being JAK1 JH2 and the other being bone morphogenic protein type II (BMPR2) receptor kinase. Its inhibition concentration *K*_i_ for TYK2 JH2 is 0.02 *n*M, which is 17-fold more selective over JAK1 JH2 and 10,000-fold over BMPR2 (Burke et al., 2019). Unexpectedly, the isolated TYK2 JH2 domain is not very stable with reported median melting temperature of only 43.5°C, which is only a few degrees above human body temperature (Min et al., 2015; Wrobleski et al., 2019). DEU and other TYK2 JH2 specific inhibitors were developed from lead compounds that were initially observed in large-scale biochemical screens of libraries of chemicals and targets (Fig. 2) (Tokarski et al., 2015). The binding of DEU and its derivative inhibitors increases the midpoint of thermal denaturation temperature to 66.0°C in the presence of 1 μM of selected inhibitors and as high as 70.2°C in the presence of 10 μM (Liu et al., 2023; Min et al., 2015). Moreover, the inhibition mechanism by DEU exhibits a large thermodynamic isotope effect between deuterium-substituted and light H methyl groups at its head (Fig. 2). Aside from these biophysical properties of these inhibitors on TYK2 JH2, little is known about their inhibitory and structural mechanisms.

**Figure 2.**
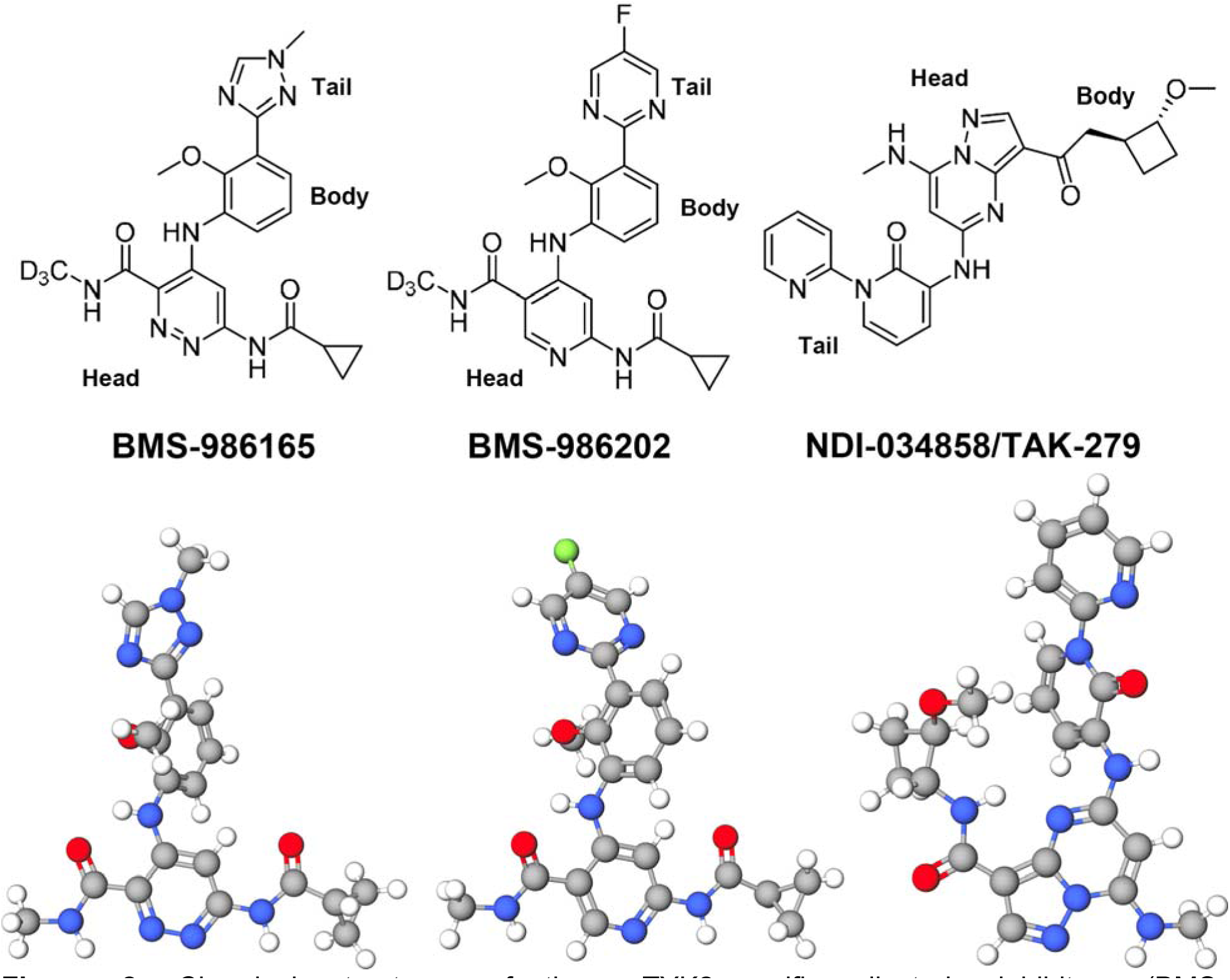
Chemical structures of three TYK2-specific allosteric inhibitors (BMS-986165/Deucravacitinib; BMS-986202; TAK-279) in advanced stages of clinic trials or Food and Drug Administration (FDA)-approved for some diseases (i.e. psoriasis) (top) and one selected conformational structure for each inhibitor (bottom).

In this study, a 3D model is provided for phosphorylation of uSTAT proteins and assembly of activated TYK2/JAK1 heterodimers based on recently reported cryo-EM structures of homodimeric (JAK1/IFNλR1)_2_. We provide computational evidence for a triple-action inhibitory mechanism of these allosteric inhibitors. Our model suggests that an inhibited state of TYK2 JH2 binds to the backside of the JH1 ATP-binding cleft and engages with the JH1 recognition loop that interacts with the adenine base of ATP. This keeps JH1 in a closed cleft conformation, making the γ-phosphate of bound ATP inaccessible to the nucleophilic hydroxyl group of the Tyr residue. To open the β-hairpin of the GXGXΦG motif (where Φ is either Phe or Tyr residue) and grant access to the γ-phosphate of ATP in JH1, the cleft must undergo a 14° rotation. The analysis of activated heterodimeric complexes of interferon receptors also shows that ATP bound to JH2 is directly hydrogen-bonded to the amino acid residue R895 of JH1 in an active state for trans phosphorylation – *i.e.*, the last step of the activation process.

## Results and Discussion

### Phosphorylation active site in the autoinhibited state of TYK2 JH2-JH1

Comparisons of the phosphorylation active site in the autoinhibited structure of TYK2 JH1 (PDB ID 1gag) with a structure containing an inhibitor (PDB ID 4oli), as well as the TYK2 JH2 structure in the presence of ATPγS (PDB ID 5c03), showed the inhibited TYK2 JH1 structure has a distinct conformation of the active site (Fig. 3, Video 1). This conformation exhibits a lobal rotation of 14° in addition to a 4 Å displacement corresponding to the opening-closing motion of the β-hairpin that controls accessibility of the γ-phosphate of ATP to the protein substrate (Lupardus et al., 2014; Min et al., 2015; Parang et al., 2001). Binding of DEU to TYK2 JH2 rigidifies the interface between JH2 and JH1, restricting the dynamics of TYK2 JH1 required for catalysis (Fig. 3D). The heavy H methyl group in DEU near this region might suppress the kinetic effect by further rigidifying this region, a hypothesis that needs to be explored by molecular dynamics simulations.

**Figure 3.**
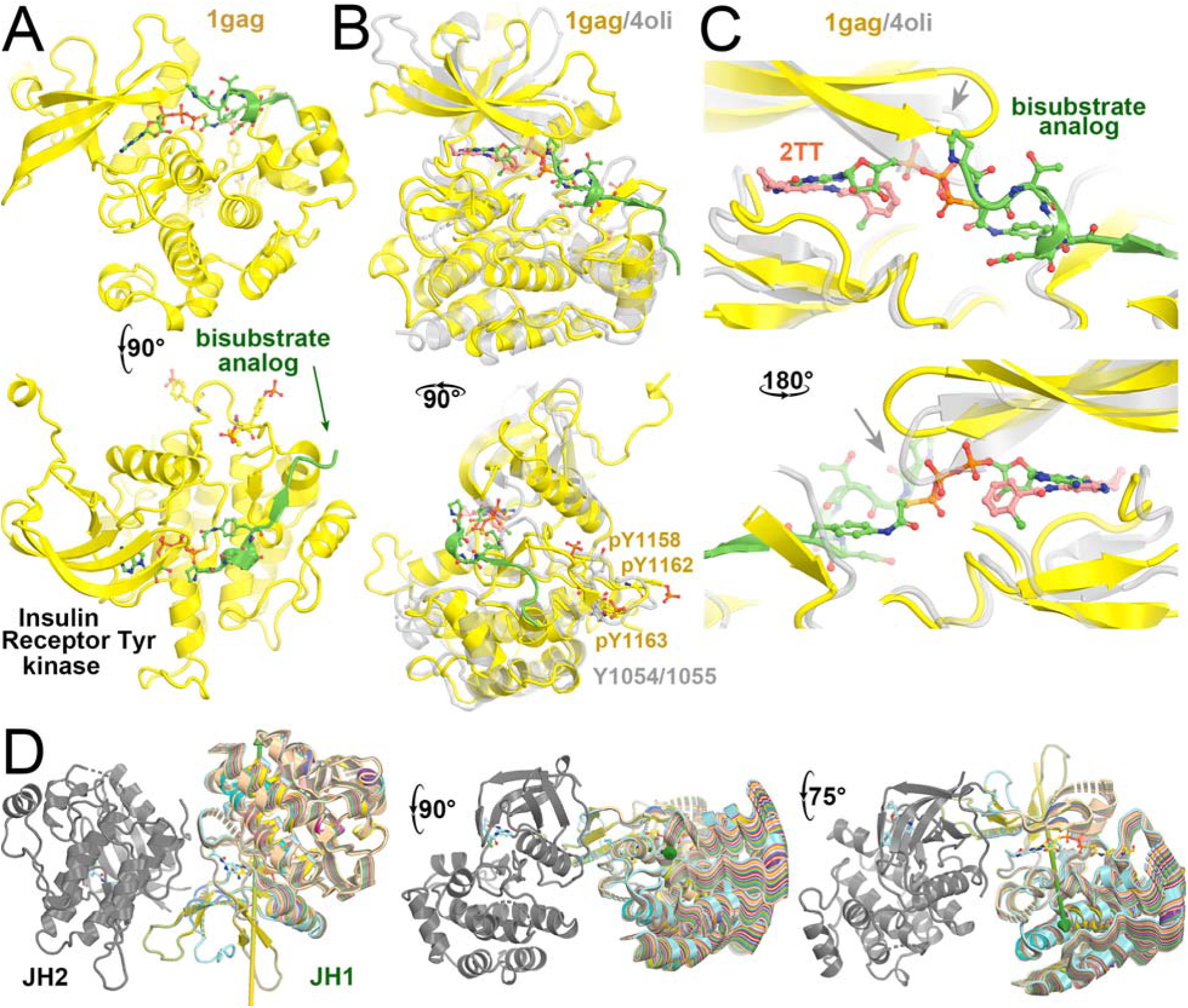
Comparison of IRTYK with TYK2 JH2 reveals subtle lobal rotations for activation. (A) Two orthogonal views of IRTYK. (B) Two orthogonal views of superposition of TYK2 JH2 (grey) with IRTYK (gold). (C) Back and front closeup views of the cleft of the two superimposed structures. Gray arrow highlights the displacement of β-hairpin as a result of subtle domain rotation. (D) Three views of predicted structural transitions of TYK2 JH2 according to comparison with IRTYK with intermediate stages of the hypothetically modeled transition in multiple colors taken from our video. See Video 1 for animation.

Evaluation of selected TYK JH2-inhibitor complexes and JH2-ATPγS complex revealed that the cleft does not exhibit significant opening and closing motions upon binding of ATPγS or inhibitors (Fig. 4), which is consistent with small-angle X-ray scattering studies (Lupardus et al., 2014; Min et al., 2015; Tokarski et al., 2015). Comparison of TYK2 JH1 in the autoinhibited complex (PDB ID 4oli) with the activated JAK1 JH1 complex (PDB ID 7t6f) showed that the cleft in the inhibited TYK2 JH1 structure is more closed than in the activated JAK1 JH1 structure. The inhibited TYK2 JH2 structure also has a cleft that is more closed than in the activated JAK1 JH2, whereas there is little difference in the extent of the closing/opening state of the clefts between TYK2 JH1 and JH2, and between JAK1 JH1 and JH2 (Fig. 5) (Glassman et al., 2022; Lupardus et al., 2014). Although the extent of closing/opening state of the cleft between JAK1 JH1 and JH2 domains remains unchanged, the entire ADP can be fitted into the cryo-EM map for JAK1 JH1. However, the orientation of the nucleotide is different in JAK1 JH2, with no visible features for its diphosphate in the cryo-EM map.

**Figure 4.**
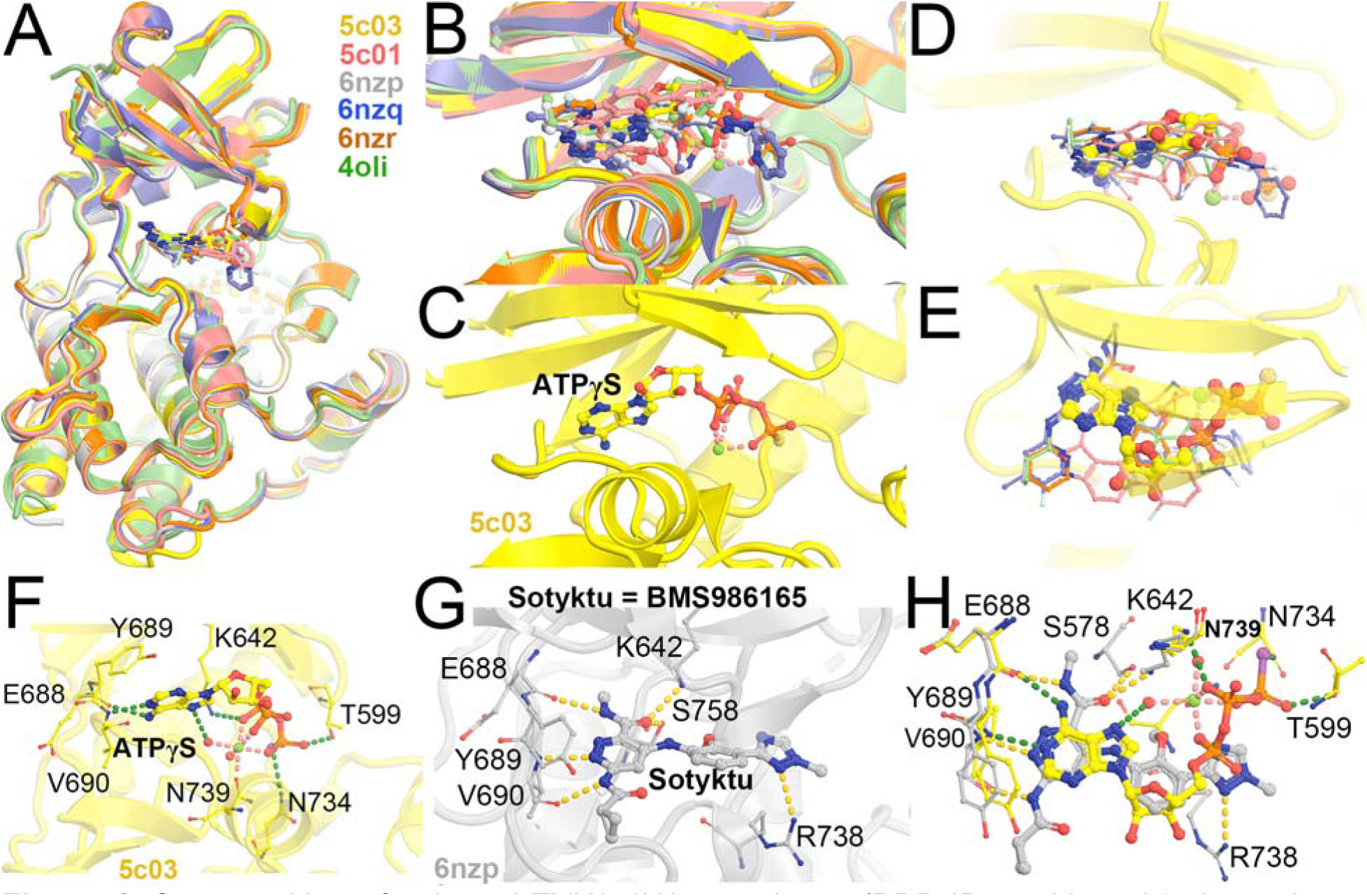
Superposition of selected TYK2 JH2 complexes (PDB IDs 5c03, 5c01, 6nzp, 6nzq, 6znr, and 4oli). (A) Overall structure. (B) Closeup view of the binding pocket. (C) ATPγS-bound 5c03 complex. (D, E) Two orthogonal views of 5c03 structure with ATPγS and DEU (its three-letter residue code 2TT as used in the PDB) from 6nzp structure. (F) Detailed hydrogen bond interactions of ATPγS with TYK2 JH2 in 5c03 structure. (G) Detailed HB interactions of DEU/Sotyktu with TYK2 JH2 in 6nzp structure. (H) A combined view of ATPγS and DEU.

**Figure 5.**
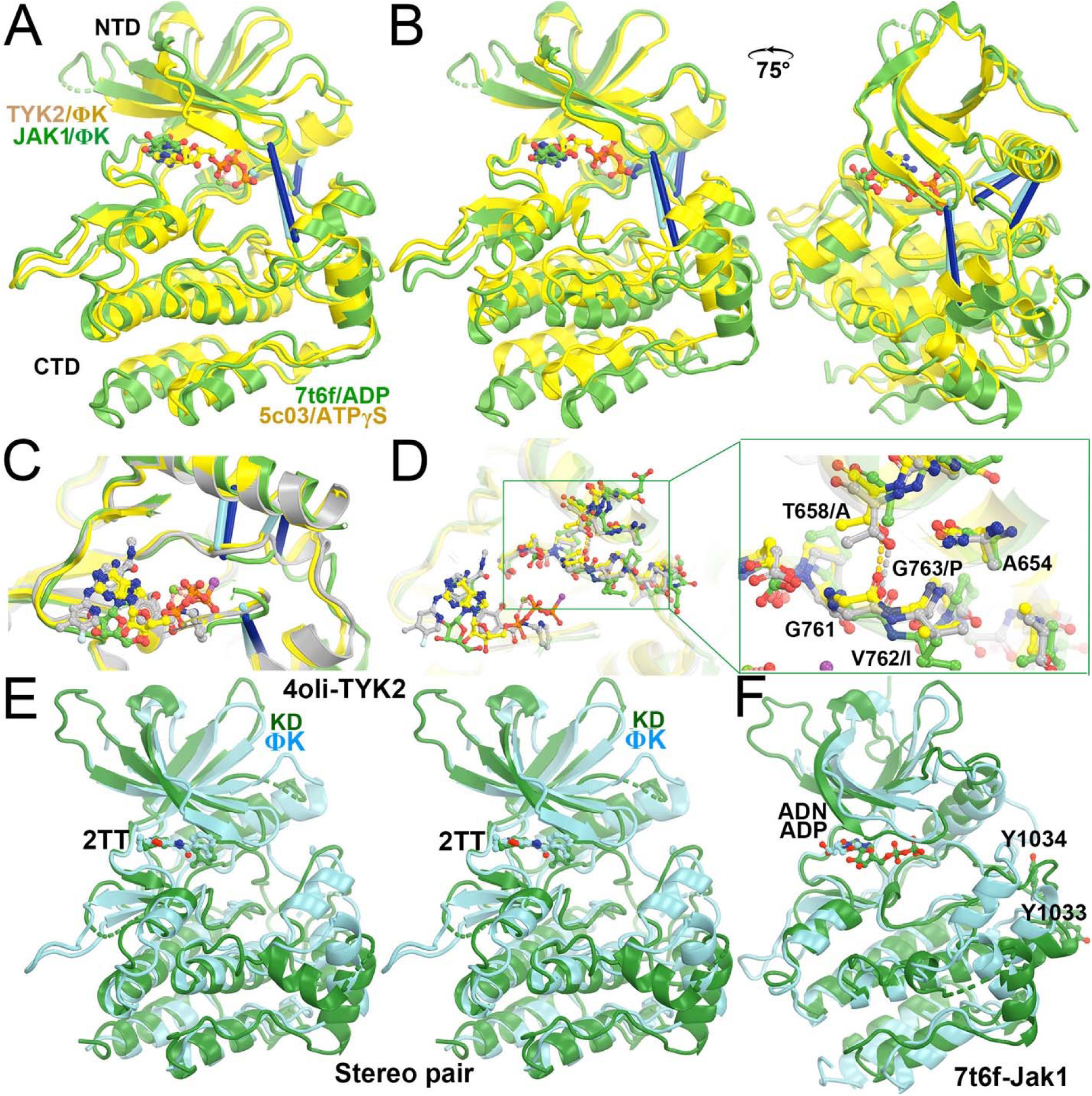
Structural basis of DEU selectivity for TYK2 over JAK1 and conformational differences between the two proteins. (A) Alignment of TYK2 JH2 (gold) and JAK1 JH2 (green) using C-terminal lobe. Three lines indicates different extents of closing-and-opening of the cleft. (B) Two views of alignment using N-terminal lobe. (C) A closeup view of (B) shows two different Cα distances. The TYK2 JH2-DEU (grey) structure is also added for comparison. (D) A further closeup of residue differences between the two JH2 domains (see explanation in text). (E) Stereodiagram of comparison of TYK2 JH2 (cyan) and JH1 (green), both with DEU bound when a very high concentration of DEU was included in crystallization. (F) Comparison of JAK1 JH2 (cyan) and JH1 (green).

The cleft of TYK2 JH2 is more closed than that of JAK1 JH2, likely due to differences in the primary sequences of the two JH2 domains since there is no evidence for inhibitor-induced changes in the cleft of TYK2 JH2 that could explain the observed differences. The G761-V-G763 motif in TYK2 JH2 is in close proximity to a nearby helix establishing backbone H bonding interactions with the T658 sidechain (Fig. 5D). In JAK1 JH2, residues of the GVG motif are instead GIP, thereby preventing tight packing in the third position. Furthermore, an Ala residue occupies the T658 position, eliminating the backbone H-bond of the first position. These differences in primary sequence explain the observed differences in the cleft and why DEU binds more tightly in the TYK2 JH2 cleft than in the cleft of JAK1 JH2 (i.e., inhibition selectivity).

We produced a full-length TYK2 model in the autoinhibited state (Glassman et al., 2022; Lupardus et al., 2014; Wallweber et al., 2014) (Fig. 6, Video 2) by aligning the TYK2 FERM-SH2 structure (PDB ID 4po6) and the autoinhibited TYK2 JH2-JH1 structure (PDB ID 4oli) onto the activated JAK1 JH1 complex model (PDB ID 7t6f). The alignment for the TYK2 JH2-JH1 (PDB ID 4oli) model used the N-terminal lobe of TYK2 JH2. We identified extensive interactions between TYK2 JH1 and TYK2 FERM-SH2, including stacking interactions between Y965 and Y376, hydrogen bonds between E957 and R381 and between K1170 and Y123, and between E1163 and the backbone carbonyls of the P116-P120 stretch (Fig. 6). Given that the N-terminal and C-terminal lobes of the two JH2 differ in orientations, an alignment using the C-terminal lobe resulted in a slight opening of the TYK2 JH1 and FERM-SH2 interface. We expect that these two alignments provide a limiting boundary of our modeled complex relative to the experimental structure yet to be determined. This interface could further reduce the dynamics of TYK2 JH1, establishing a target for a new kind of inhibitors.

**Figure 6.**
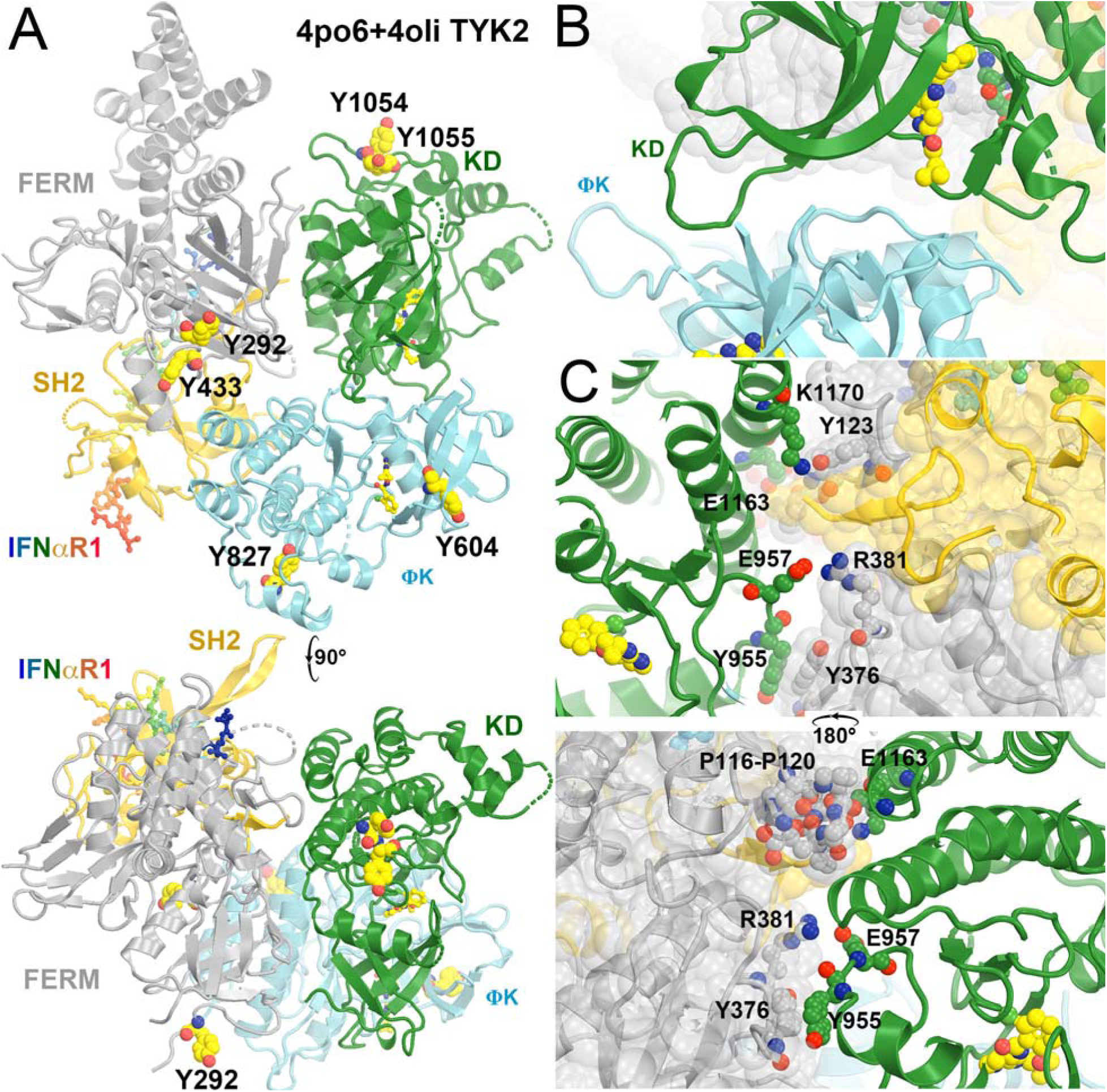
Modeled full-length TYK2 in the autoinhibited state in complex with IFNαR1. (A) Two orthogonal views. Locations of pY residues upon activation are large balls-and-sticks models. (B) A closeup view of the JH2-JH1 interface. (C) Two back and front closeup views of the FERM-SH2/JH1 interface. See Video 2 for animation.

### Two active states of the JAK1/IFN**λ**R1 complex for modeling TYK2/JAK1

We modeled two activated states of the heterodimeric TYK2/JAK1 complexes using two homodimeric (JAK1/IFNλR1)_2_ complexes (PDB ID 7t6f and 8ewy) (Figs. 7, 8, Videos 3, 4). Equivalent TYK2 models have been previously generated using AlphaFold (Caveney et al., 2023). Below, we provide new mechanistic insights into the JAK-STAT signaling pathway and the inhibitory mechanism by DEU upon binding to TYK2 JH2.

**Figure 7.**
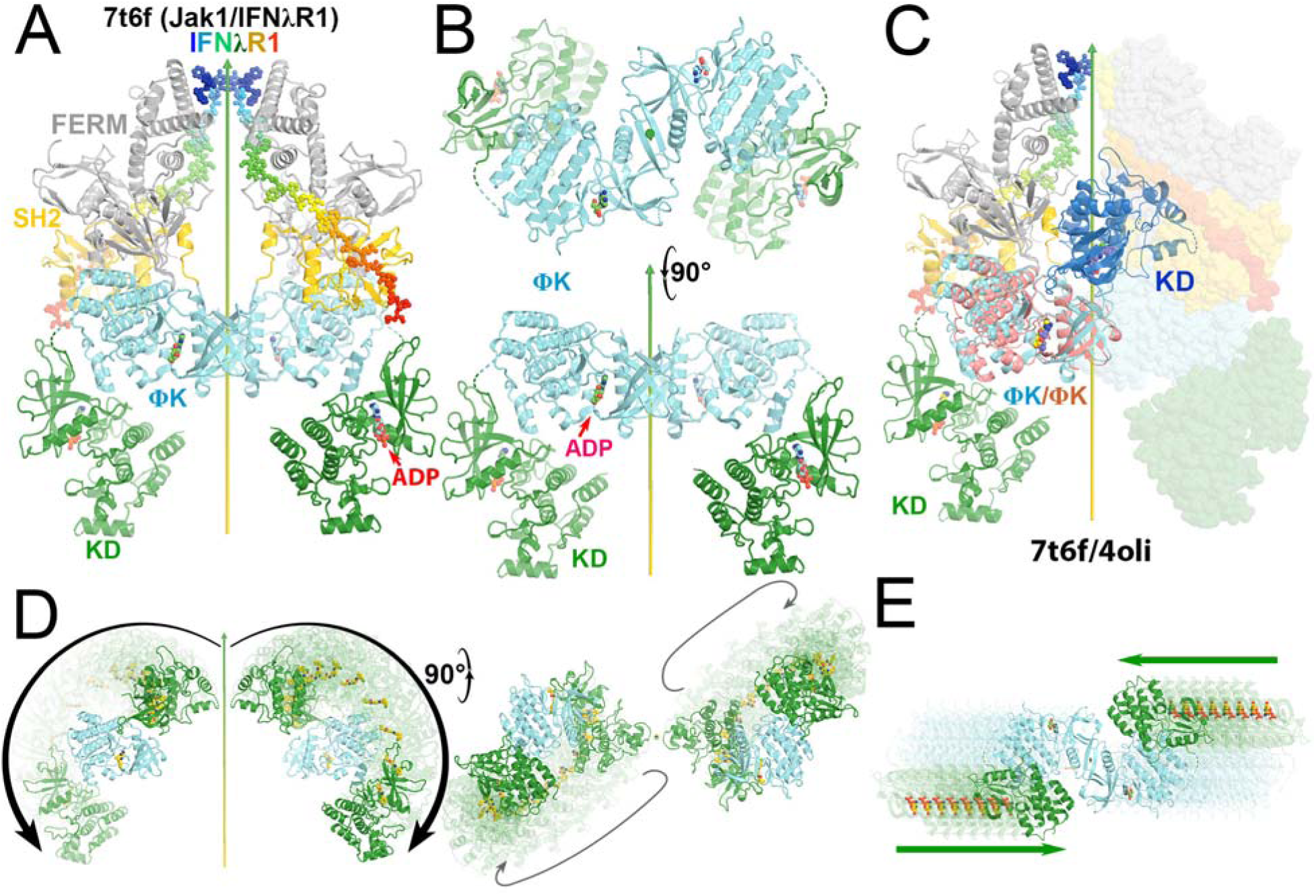
Comparison of the autoinhibited and the active state for downstream phosphorylation of uSTAT substrates of JAK1 and TYK2. (A) The full-length (JAK1/IFNλR1)_2_ complex colored by domain. The IFNλR1 receptor are shown in rainbow balls-and-sticks model. (B) Two orthogonal views of JH2-JH1 domain in the active state. (C) Alignment of one autoinhibited TYK2 JH2 (salmon)-JH1 (blue) domains onto the JAK1 complex with one subunit in ribbons and the other in surface model. (D) Two orthogonal views of proposed domain rotation morphing intermediate stages of the hypothetically modeled transition taken from our video. (E) Proposed domain/subunit translation taken from video. See Video 3 for animation.

**Figure 8.**
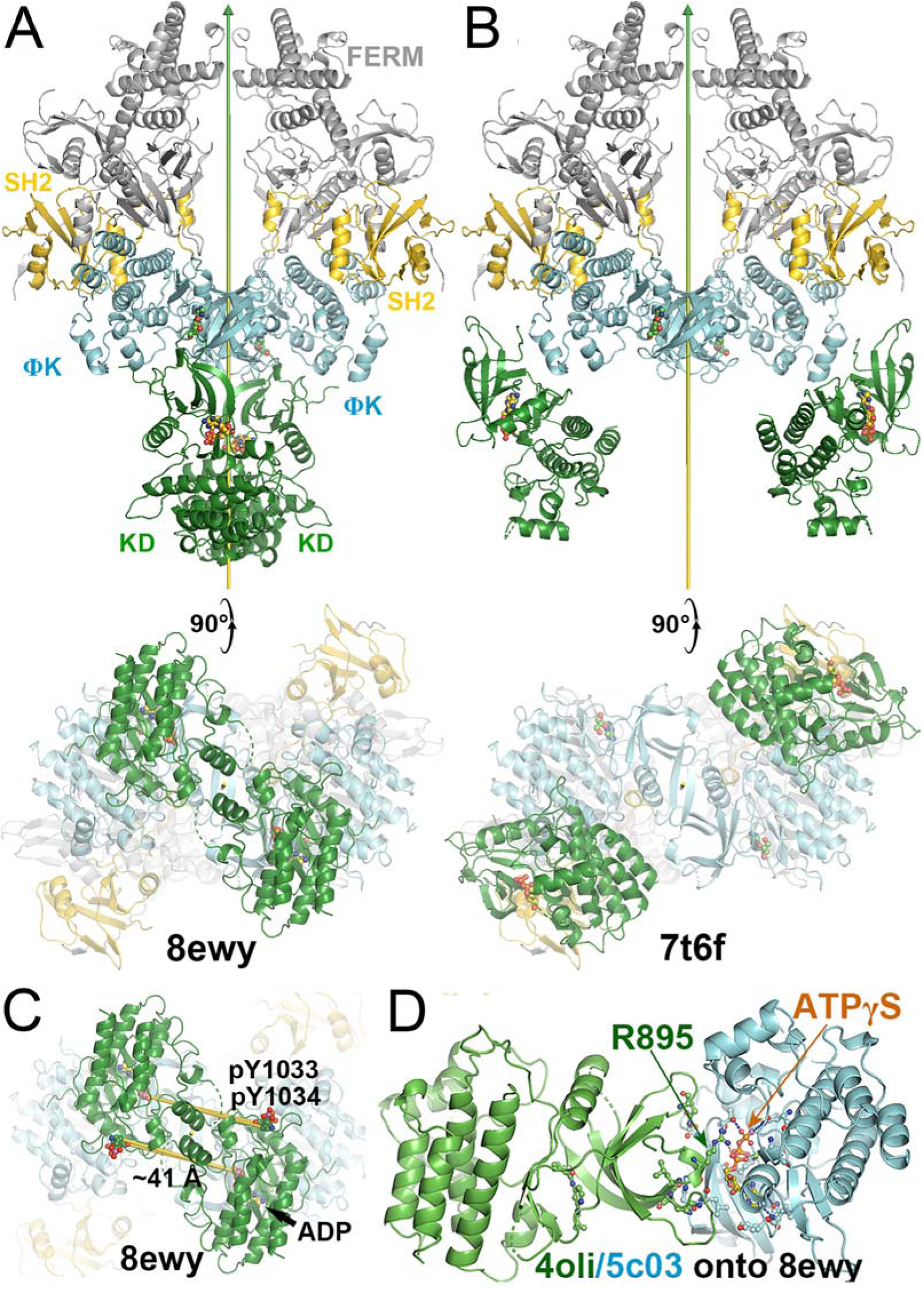
Comparison of two active states of the (JAK1/IFNλR1)_2_ complexes and conformational conversion between them. (A) Two orthogonal views of the active state for trans phosphorylation colored by domains. (B) Two orthogonal views of the active state for downstream uSTAT phosphorylation. (C) Distances of trans pY sites to the catalytic site of their partner, indicating additional conformational changes of substrate loop is required for pYpY sites to be phosphorylated. (D) A closeup of the JH2-JH1 interface for modeled TYK2 protein including an ATPγS bound in the TYK2 JH2. See Video 4 for animation.

These two active states of the heterodimeric TYK2/JAK1 complexes exhibit two distinct modes of interaction between TYK2 JH2 and JH1, each with unique interfaces. These differ substantially from the interaction mode of the autoinhibited state with different interfaces. In reference to TYK2 JH2, we find a domain rotation of TYK2 JH1 by 230° (but not 130° = 360° - 230°, see below for rotation direction) between the autoinhibited (PDB ID 4oli) state and the first (PDB ID 7t6f) active state (Video 3). It is also 230° (but not 130° = 360° - 230°) between the first (PDB ID 7t6f) and the second (PDB ID 8ewy) active states (Video 4). Coincidentally, it is also 230° (but not 130° = 360° - 230°) between the autoinhibited (PDB ID 4oli) and the second (PDB ID 8ewy) active states. However, rotations occur along different axes nearly tangentially on the TYK2 JH2 surface, *i.e.*, JH1 approximately rolls over the JH2 surface. Although rotation angles are approximately the same in transitions between any two of the three states, the centers of interacting surface of JH2 with JH1 between the two active states are closer than those between the autoinhibited state and one of the two active sites in isosceles triangular relationship on JH2 surface (Videos 3, 4). In the second (PDB ID 8ewy) active state, two JH1 domains of the two enzymes within the activated dimer also interact directly with each other.

Energetically, interactions between TYK2 JH2 and JH1 in the autoinhibited (PDB ID 4oli) state should be far stronger than both active states (PDB IDs 7t6f and 8ewy), as shown by experiments (Lupardus et al., 2014). In fact, when the domains of TYK2 JH2 and TYK2 JH1 were mixed, the phosphorylation activity of TYK2 JH1 was reduced by 2 orders of magnitudes (Lupardus et al., 2014), suggesting that only the autoinhibited state of the formed TYK2 JH2-JH1 complex was present in the reaction mixture. In the full-length TYK2 model, additional interactions were observed between the TYK2 FERM-SH2 and JH1 domain, further stabilizing the autoinhibited state over two other activated states. These observations rule out the hypothesis that monomeric TYK2 establishes an equilibrium between the autoinhibited state and activated states, where the cytokine receptor dimerization simply shifts this equilibrium (Glassman et al., 2022). Such hypothesis is not supported by existing TYK2 data.

Based on our structural analysis, we propose a Velcro zipper mechanism for activation of JAKs. When IFNαR1 binds TYK2 FERM-SH2 and IFNβR1 binds JAK1 FERM-SH2, dimerization of the two receptors brings the TYK2 and JAK1 monomers closer to form a heterodimeric complex, which actively pushes the two JH1 domains of the two monomers away from each other. Therefore, JH1 has to rotate away from the central axis (i.e., 230°), without crossing over the central axis (130°). Once JH1 exits from the autoinhibited state, it becomes active for tyrosine phosphorylation. When JH1 rolls over the surface of JH2, as it transitions from the autoinhibited state to the activated state, its active site interacts with 4 specific tyrosine residues on its pathway for phosphorylation by TYK2, but none by JAK1. This phosphorylation is highly efficient because the phosphorylation is a zero-order reaction. The resulting phosphorylation differs from the *in vitro* first-order phosphorylation reaction based an isolated JH1 domain, which is linearly proportional to substrate concentration and requires a very high concentration to observe any detectable reaction. Such high concentrations of protein substrate(s) as used *in vitro* studies are only mechanistically important but not biologically relevant. Once JH1 rotates away from the central axis, two JH2 domains move closer to each other by a displacement of 35 Å for JH2-JH2 dimerization. While rotation and translation were modeled independently in our videos, the two motions likely occur simultaneously since they are quite independent from each other (as other motions discussed below). The proposed zipper mechanism also brings two pSTAT monomers together before they leave from the activated TYK2/JAK1 dimer, i.e., the activated complex also serves as a chaperone for parallel pSTAT dimerization after it completes phosphorylation of its SH2 pY residue (see below).

The teeth of Velcro zipper would need some flexibility for breathing motions for it to work properly. The binding affinity of ATP to TYK2 JH2 is at 24 μM (Min et al., 2015). Binding and transient release of ATP provide such required breathing motions. These motions are centered on the TYK2 JH2 (M978)-EYVF motif, which interacts with both the adenine base of bound ATP (or the head of bound DEU) and thereby TYK2 JH1 is in the autoinhibited state. However, the binding of DEU to TYK2 JH2 is on the order of *K*_i_ = 0.02 *n*M (Burke et al., 2019). This binding interaction suppresses the breathing motion; therefore, impeding protein dynamics is part one of allosteric inhibitory mechanism. Replacement of the heavier trideuterium methyl group by a lighter H methyl group or N relative to C atom at a head ring of DEU further increases the breathing motions of both the inhibitor and this interacting loop (Wrobleski et al., 2019), and thereby reduces inhibition.

TYK2 JH1 modeled in the second active state (PDB ID 8ewy) makes extensive interactions with JH2, centered at the bound ATPγS to the JH2 domain when the TYK2 JH2-ATPγS complex (PDB ID 5c03) was aligned onto the 8ewy complex (Fig. 9A, 9B) (Caveney et al., 2023; Min et al., 2015). This active complex is essential for trans phosphorylation of tyrosine residues in JH1 by its partner’s JH1 kinase domain for geometric reasons, i.e., the TYK2 JH1 pY1054/pY1055 doublet sites by JAK1 JH1, and the JAK1 JH1 pY1033/pY1034 doublet sites plus its pY1125 site by TYK2 JH1. Without any optimization in our model, R895 of JH1 binds the α-phosphate group of ATPγS bound to JH2 plus there are four other potential inter-subunit H bonds: (i) Q969 of TYK2 JH1 with T599 of TYK2 JH1, (ii) R895 of JH1 with backbone amide of L595 (and perhaps of H594), and (iii) L897 backbone carbonyl with the H594 sidechain (Fig. 9, Video 5). In addition, K894 next to R895 can enhance electrostatic interactions with the triphosphate moiety of ATP. Results of this model are fully consistent with those of a full-length TYK2 K642A mutant in its JH2, which eliminated its basal ATP hydrolysis bound to JH2 but retained some ATP binding ability (Min et al., 2015). Therefore, ATP binding to TYK2 JH2, but not its ATP hydrolysis, is essential for the full activation of this kinase. DEU has no equivalent trisphosphate moiety of ATP in its tail. Binding of DEU to TYK2 JH2 competitively inhibits ATP binding in TYK2 JH1, which is part two of the allosteric inhibitory mechanism.

**Figure 9.**
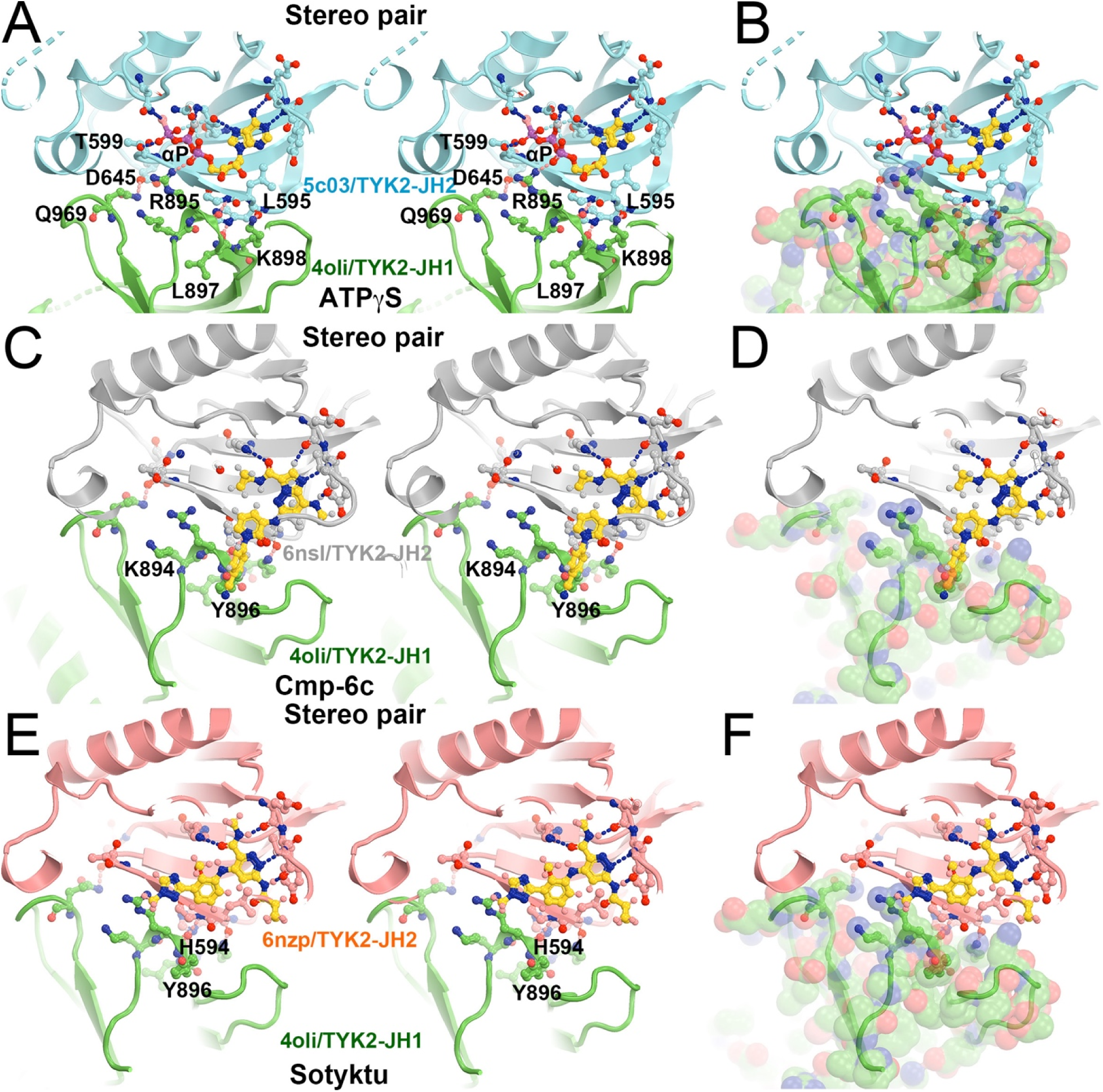
Roles of ATP and TYK2-specific inhibitors during trans phosphorylation. (A, B) Stereodiagram of selected residues (A) and view of the full-surface representation of TYK2 JH1 (B) in the interface between TYK2 JH2 (cyan) and TYK2 JH1 (green) mediated by ATPγS. (C, D) The KZJ (or compound C)-inhibited complex of TYK2. (E, F) The DEU/Sotyktu-inhibited complex of TYK2. See Video 5 for animation.

When compound KZJ in the TYK2 JH2 complex (PDB ID 6nsl) was modeled in the second active state (PDB ID 8ewy) (Caveney et al., 2023; Liu et al., 2023), its cyanophenyl group completely overlaps with the Y896 sidechain of TYK2 JH1 (Fig. 9C, 9D). KZJ is a member of imidazo[1,2-b]pyridzine derivatives, which differs from the chemical backbone structure of DEU (Figs. 2, 10). This modeling shows that this compound not only competitively inhibits ATP binding, but also directly interferes with JH1 approaching JH2 in this position. Our modeling further shows that NDI-034858 and KZJ belong to the same structural group and exhibit the same inhibitory mechanism although the chemical backbone structures of these compound differ (Fig. 10). Although the tails of these compounds do not contribute much to their binding to TYK2 JH2 because the tails stick out to the surface of JH2 but are located at the domain interface in this modeled complex, these tail protrusions provide a structural basis for prevention of formation of this active complex. Similarly, the tail of DEU also has an overlapping problem with the R895 sidechain and should have a similar structural property. Thus, steric clashes of DEU with JH1 residues form part three of the triple-action inhibitory mechanism of these compounds. Our modeling differs from an early attempt of AlphaFold-based modeling (Caveney et al., 2023), which failed to explain how DEU allosterically inhibits the activation of the TYK2/JAK1 heterodimer because they were trying to optimize the interactions that DEU actually destabilizes between the TYK2 JH2 and JH1 domains.

**Figure 10.**
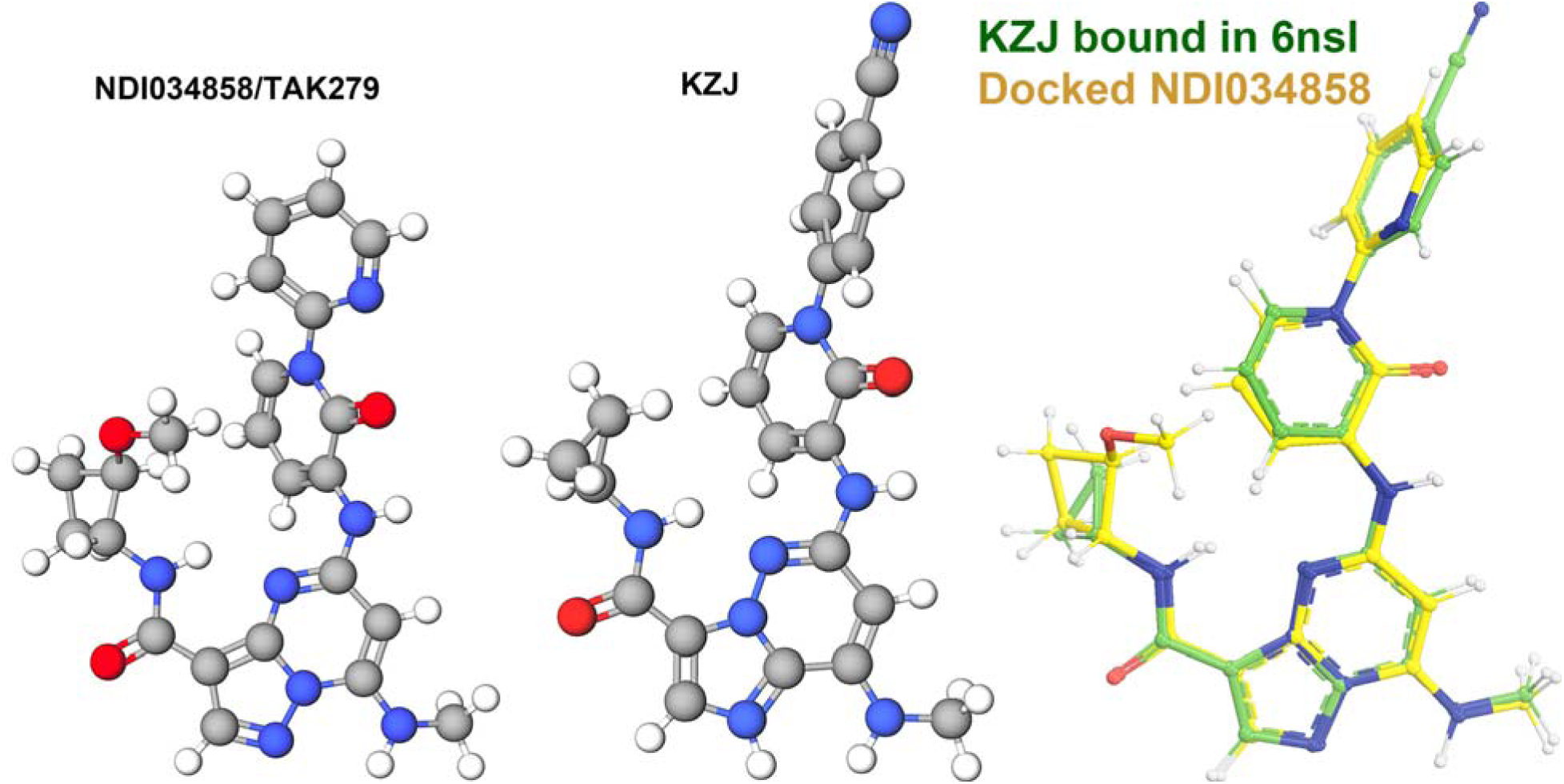
Modeling of NDI-034858/TAK279 (left) using compound KZJ (center) shows that they are structurally related compounds (right) even though their chemical background structures differ.

### The activated TYK2/JAK1 heterodimeric complex as a streamline factory

To understand how the activated TYK2/JAK1 heterodimeric complex recruits stable antiparallel uSTAT1 dimers and uSTAT2 dimers, phosphorylates them, and assembles new stable parallel pSTAT1 dimers and pSTAT1-pSTAT2 heterodimers as products, we modeled a TYK2/JAK1 heterodimeric complex using the (JAK1/IFNλR1)_2_ homodimer by replacing one copy of JAK1 with TYK2 and visualized the distribution of pY residues on the complex. JAK1 has three pY sites, doublet pY1033/pY1034 and pY1125, all in JH1. However, our examination of the cryo-EM atomic model (PDB ID 7t6f) showed that pY1125 actually corresponded to Y1124 in the coordinate file with one residue shift in this region for an unknown reason. TYK2 has 6 pY sites, pY292, pY433, pY604, pY827, and doublet pY1054/pY1055. We found that an alternative rotamer was required to rotate the Y827 sidechain from a buried position to an exposed position to be phosphorylated.

An examination of pY sites on TYK2 FERM-SH2 and JH2 shows that they are on the rotational path of TYK2 JH1 that we proposed above between the autoinhibited state and the first activated state (PDB ID 7t6f) for cis autophosphorylation, and that none are on the path of the opposite rotational direction. How different JAK proteins have different patterns of pY sites may depend on specific cytokine receptors within the activated complexes. TYK2 JH1 cannot auto phosphorylate its own pY1054/pY1055 doublet site due to geometric restraints, which have to be phosphorylated in trans by JH1 of its dimeric partner, i.e., JAK1 JH1 in our modelled complex. While TYK2 JH1 has only the pY1054/pY1055 doublet site, JAK1 JH1 has three pY sites, JAK2 JH1 has four pY sites, and JAK3 JH1 has a different set of four pY sites (Fig. 1). How these different sites are selected for trans phosphorylation remains unclear. Within 4.5 Å of the pY1054/pY1055 doublet site of TYK2 JH1, there is another pair of exposed Y1079/Y1080 site that are not known to be phosphorylated. Moreover, Y1013, 1019, and 1076 of TYK2 JH1 nearby are not phosphorylated, either. Clearly, the selection of Tyr residues for trans phosphorylation is highly specific and the understanding of this selection remains a mechanistic challenge. It is likely that our modeled TYK2 JH2-JH1 interface in the active state for trans phosphorylation may be more specific for TYK2 than for the other three JAKs where some rotation of this interface may be permissive for other pY sites to be phosphorylated besides the pY/pY doublet site. If this indeed is true, all TYK2-specific allosteric inhibitors will not have significant effects on the three other JAKs in this step of activation.

All pY sites have a potential to serve as recruitment stations for binding of uSTATs via docking into the vacant pY-binding pocket of their SH2 because these Tyr residues must be exposed and protruded away from its own protein surface for access to the γ-phosphate of ATP bound to JH1 and because domain size of uSTAT-SH2 is similar to that of JH1 (Fig. 11). However, the surface curvatures of SH2 and JH1 for binding of this tyrosine residue are not exactly the same so that different pY sites may exhibit different affinities. Given the observations that all pSTAT dimers are parallel (Becker et al., 1998; Chen et al., 1998; Li et al., 2016) and that all uSTAT dimers (PDB ID 1yvl/uSTAT1, PDB ID 1y1u/uSTAT5A, PDB ID 6tlc/uSTAT3) are antiparallel, all of which are superimposed very well within each state (Fig. 11C, Table 1) (La Sala et al., 2020; Mao et al., 2005; Neculai et al., 2005), all JAKs should be activated in similar manners to some extent. We are aware of that some higher order assemblies of JAKs are known to exist (Spinelli et al., 2021), which would require additional modeling.

**Figure 11.**
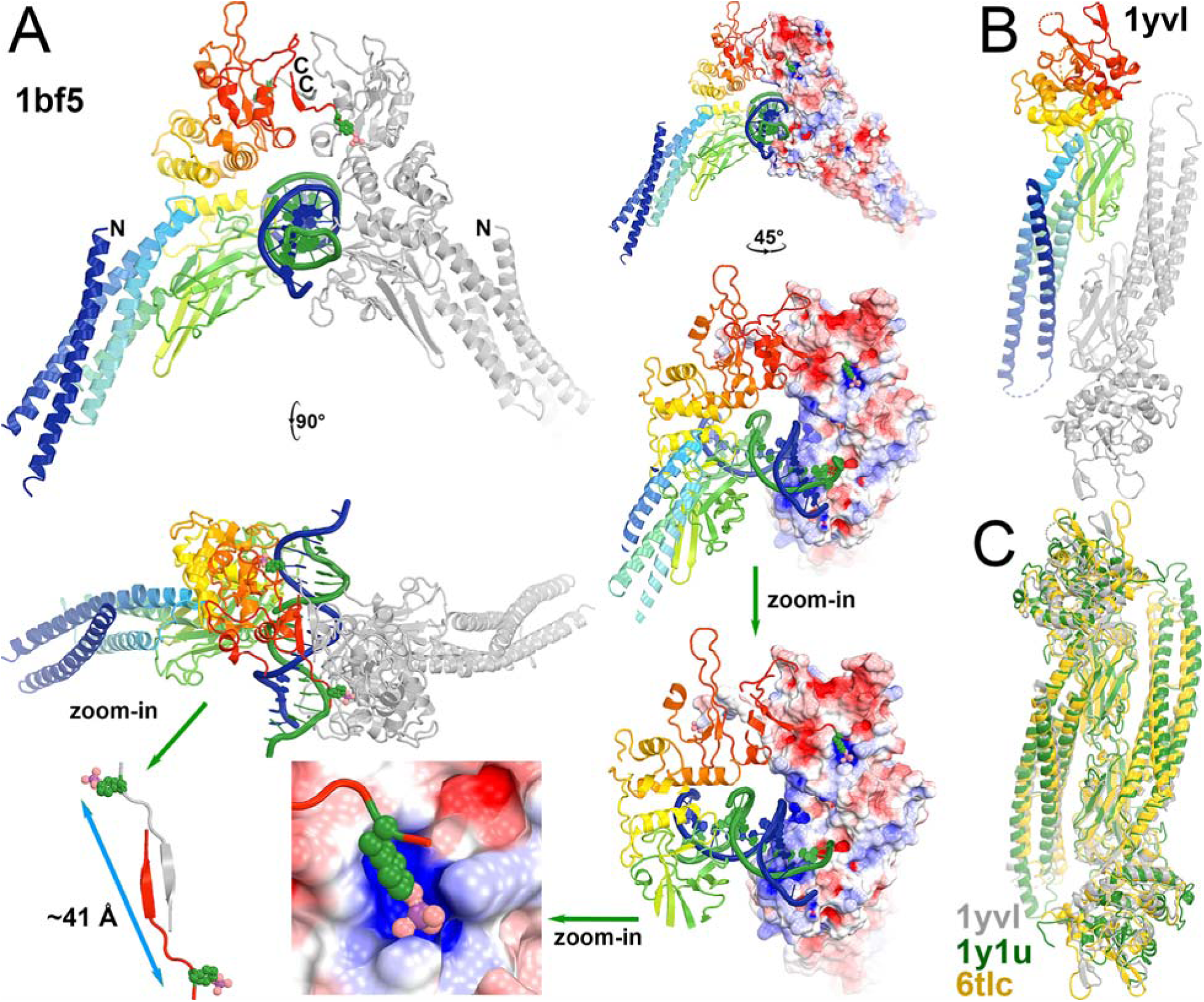
The product parallel pSTAT1 dimer and the substrate antiparallel uSTAT dimers of the activated TYK2/JAK1 heterodimeric complex. (A) Multiple closeup views of phosphorylation-dependent SH2 dimerization of the pSTAT1 dimer and details of binding pY residue into the pY-binding pocket of neighboring subunit as the means of docking of TYK2 pY onto the pY-binding pocket of SH2 in uSTAT. (B) Anti-parallel uSTAT1 dimer. (C) Conserved antiparallel uSTAT dimers of uSTAT1 (PDB ID 1yvl, grey), uSTAT5a (PDB ID 1y1u, green), and uSTAT3 (PDB ID 6tlc, gold) as a basis of docking onto the TYK2/JAK1 complex.

**Table 1.**
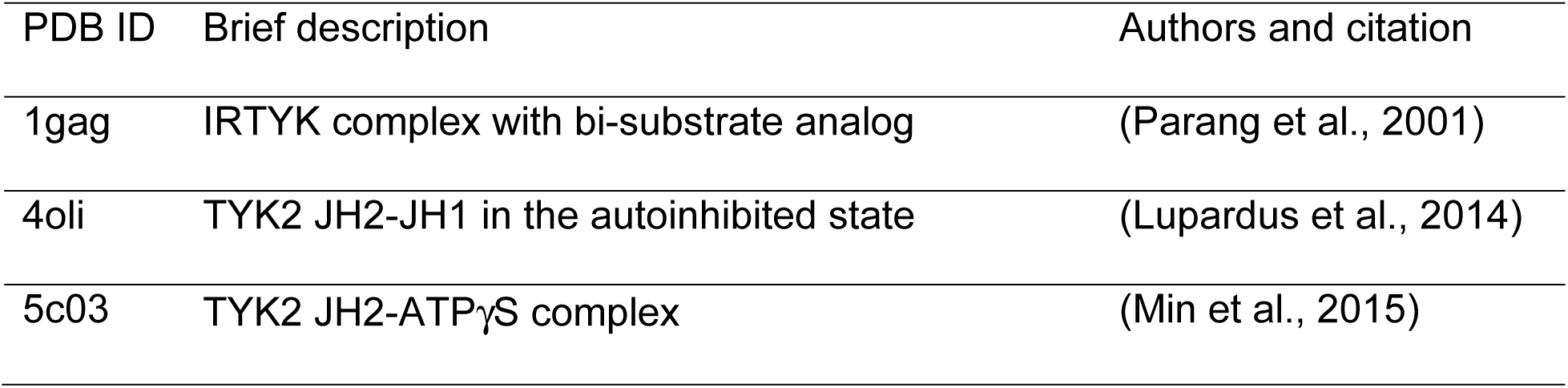
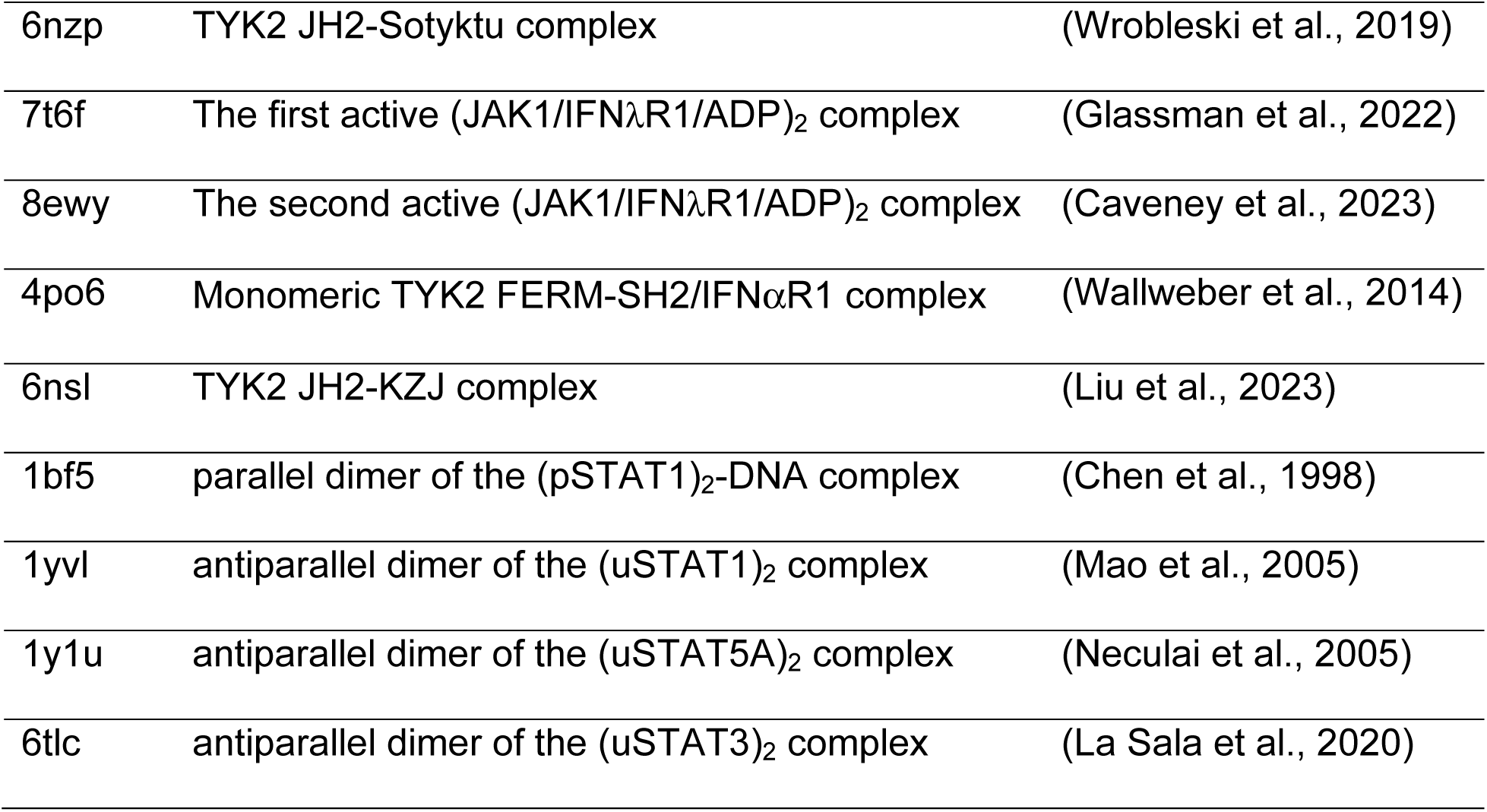
key structural information references.

| PDB ID | Brief description | Authors and citation |
| --- | --- | --- |
| 1gag | IRTYK complex with bi-substrate analog | (Parang et al., 2001) |
| 4oli | TYK2 JH2-JH1 in the autoinhibited state | (Lupardus et al., 2014) |
| 5c03 | TYK2 JH2-ATPγS complex | (Min et al., 2015) |
| 6nzp | TYK2 JH2-Sotyktu complex | (Wroblewski et al., 2019) |
| 7t6f | The first active (JAK1/IFNλR1/ADP)<sub>2</sub> complex | (Glassman et al., 2022) |
| 8ewy | The second active (JAK1/IFNλR1/ADP)<sub>2</sub> complex | (Caveney et al., 2023) |
| 4po6 | Monomeric TYK2 FERM-SH2/IFNαR1 complex | (Wallweber et al., 2014) |
| 6nsl | TYK2 JH2-KZJ complex | (Liu et al., 2023) |
| 1bf5 | parallel dimer of the (pSTAT1)<sub>2</sub>-DNA complex | (Chen et al., 1998) |
| 1yvl | antiparallel dimer of the (uSTAT1)<sub>2</sub> complex | (Mao et al., 2005) |
| 1y1u | antiparallel dimer of the (uSTAT5A)<sub>2</sub> complex | (Neculai et al., 2005) |
| 6tlc | antiparallel dimer of the (uSTAT3)<sub>2</sub> complex | (La Sala et al., 2020) |

We generated a consensus uSTAT1 dimeric model with a pY-containing peptide taken from the pSTAT1 dimer and approximately aligned this pY residue onto a target pY site of TYK2 or JAK1 while avoiding any physical overlap between the two proteins for docking. We find that multiple orientations of the uSTAT dimer can be docked onto the pY292 site of TYK2 with some rotation freedom along the pY axis (Fig. 12, Video 6). Similarly, uSTAT dimers can be docked on the second pY of the pYpY doublet site of both TYK2 and JAK1 JH1 (Fig. 12, Video 6). Furthermore, uSTAT1 dimer docked on the pY292 site of TYK2 does not overlap with one on the pY1055 site, suggesting two uSTAT1 dimers can be simultaneously present on these two sites of activated TYK2. This highlights differences between JAK1 and TYK2. Based on known distributions of pY residues on the surface of the models, TYK2 can recruit multiple uSTAT dimers to be waiting-in-line queued for rapid phosphorylation, but JAK1 cannot. Other pY sites of TYK2 between pY292 and pY1055 could provide negative electrostatic potentials and serve as intermediate docking sites as a conveyer belt for the uSTAT1 dimer bound to the pY292 site of TYK2 to move continuously onto the pY1055 site once the latter site becomes vacated. The distance between the SH2 pY binding pocket of one STAT1 and the pY701 site of its dimeric partner within the assembled pSTAT1 is about 41 Å (Fig. 11A). Distances of the pY/pY doublet sites to the catalytic site of the same subunit are about only 20 to 31 Å. When the monomer uSTAT1 was docked onto the second pY of the pY/pY doublet sites in the first active state (PDB ID 7t6f), its SH2 Y701 is within a distance to the catalytic site for phosphorylation (Fig. 13, Video 7).

**Figure 12.**
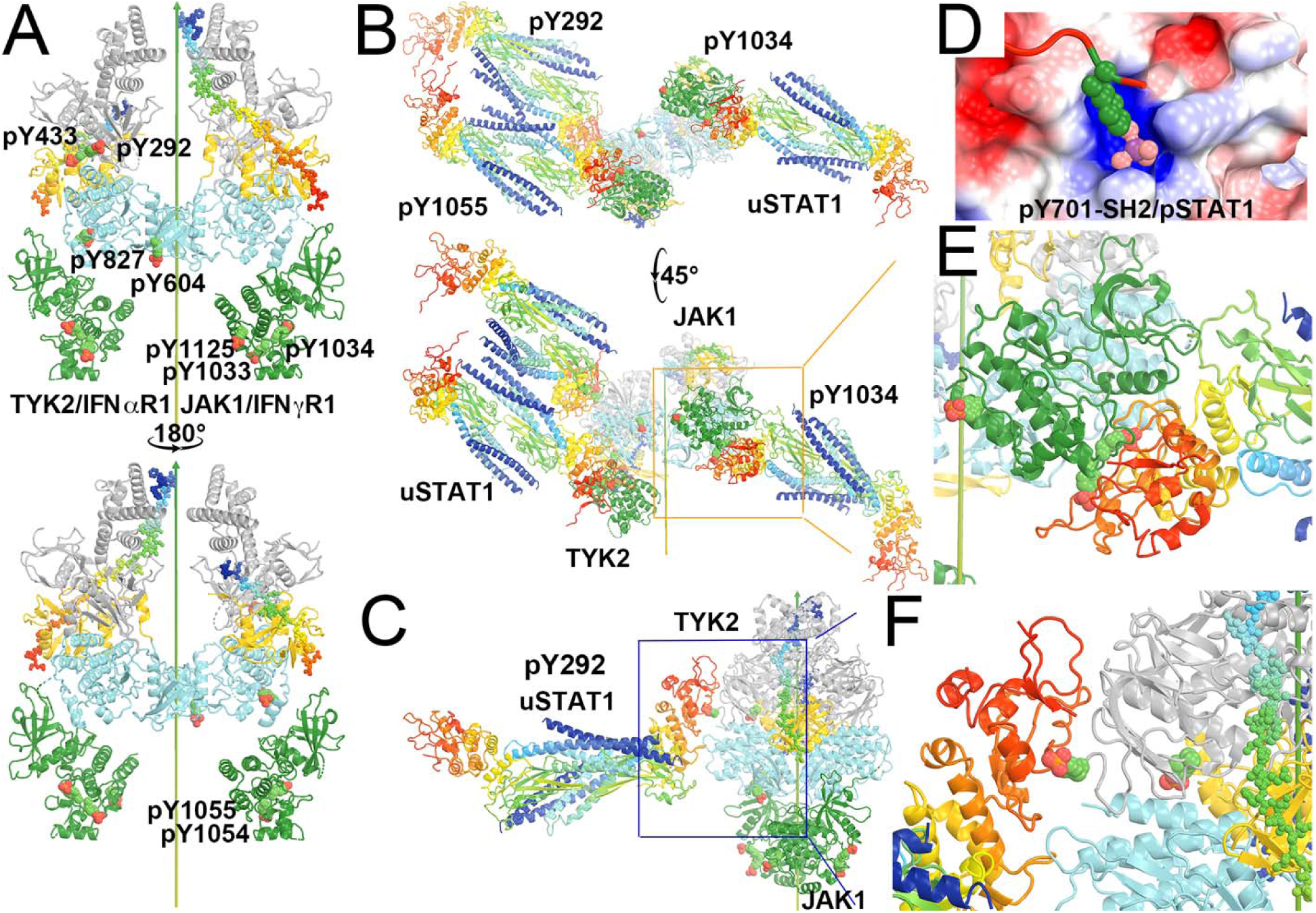
Locations of pY on the modeled TYK2/IFNαR1-JAK1/IFNγR1 heterodimeric complex where IFNγR1 represents the biologically relevant IFNβR1 and docked antiparallel uSTAT1 on selected pY sites. (A) Front and back views of the modeled complex. TYK2 has 6 pY sites (292, 433, 604, 827, 1054 and 1055) and JAK1 has three pY sites (1033, 1034, and 1125). (B) Bottom-up view and a 45° rotated view of the docked antiparallel uSTAT1 dimers on pY292 and pY1055 of TYK2 and on pY1034 of JAK1. (C) A closeup of the pY292 site of TYK2. (D) The method for docking pY onto its binding pocket according to the parallel pSTAT1 crystal structure. (E) A closeup of the pY1034 site of JAK1. (F) A further closeup of the pY292 site of TYK2. See Video 6 for animation.

**Figure 13.**
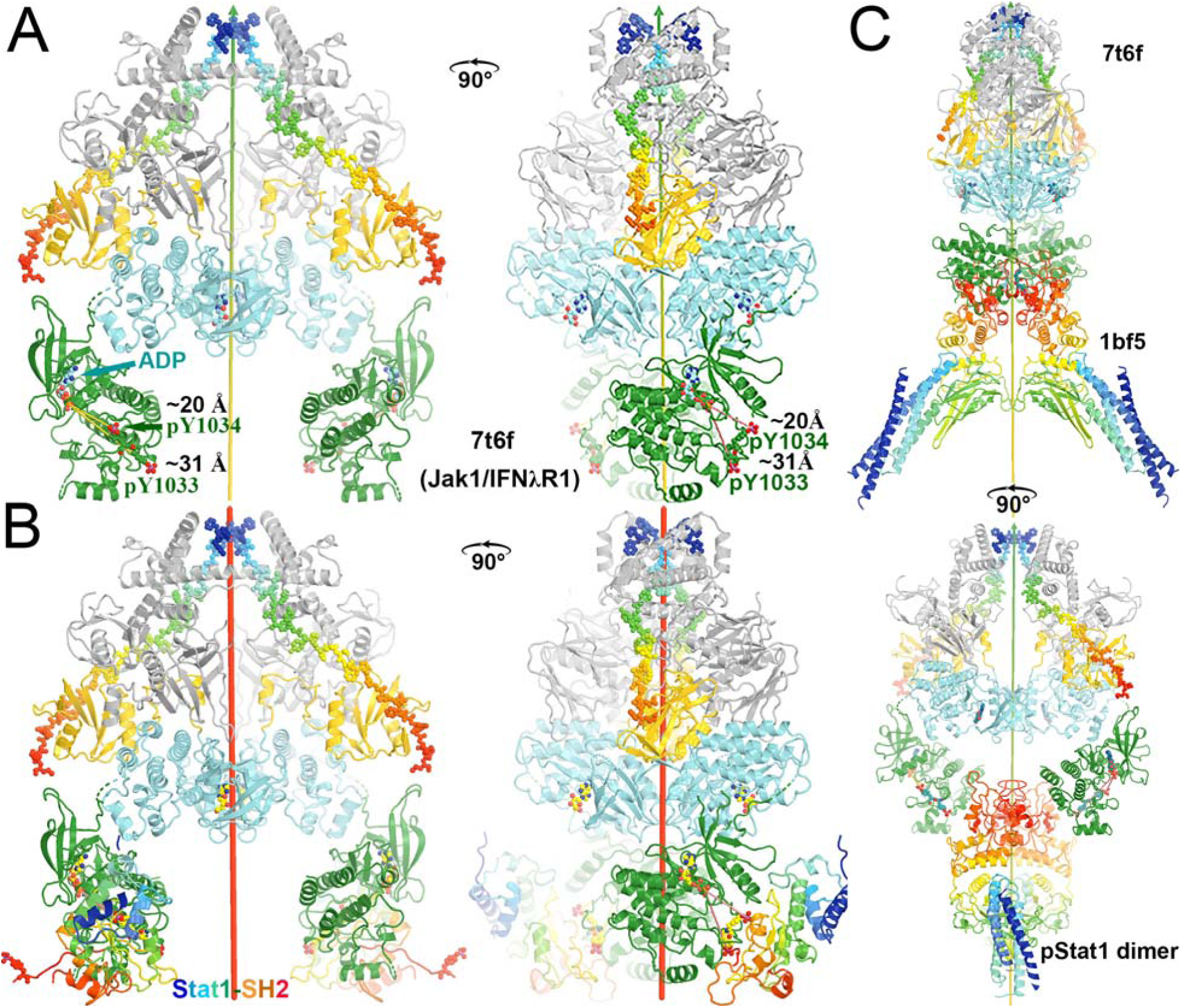
Docking of the SH2 domain of uSTAT1 onto the active state of the (JAK1/IFNλR1)_2_ complex for modeling of the active TYK2-IFNαR1/JAK1-IFNβR1 complex. (A) Two orthogonal views of the active complex colored by domains. (B) Two orthogonal views of the complex with docked SH2 domains in rainbow colors. (C) Two orthogonal views of the product parallel pSTAT1 just being assembled by the complex. See Video 7 for animation.

There were a number of asymmetric heterodimers observed whose conformations are closely related to the second symmetric active complex (Caveney et al., 2023). Functional roles of these asymmetric complexes remain unknown. We propose that they may help to transfer uSTAT substrates captured in TYK2 to its partner JAK1, which will increase the efficiency of tyrosine phosphorylation. This is particularly useful for the TYK2/JAK1 heterodimer where TYK2 captures far more queued uSTAT proteins than JAK1 can. If TYK2/IFNαR1 is specialized to capture uSTAT1 and JAK1/IFNβR1 to capture uSTAT2, the ratio of the pSTAT1-pSTAT1 homodimer and pSTAT1-pSTAT2 heterodimer can be adjusted through an inter-subunit substrate transfer mechanism. In addition, an inter-subunit product transfer mechanism may also exist, which could be useful if the activated TYK2 is primarily responsible for all pSTAT1 production, and JAK1 is primarily responsible for phosphorylation of uSTAT2 but also would serve as a storage station for some pSTAT1. This step could convert the antiparallel uSTAT1 dimer captured to the parallel pSTAT1 dimer before the dimeric pSTAT1 dimer is released. These physical properties and the proposed mechanism would rapidly collect all uSTAT dimers onto the activated complex, enabling fast responses in gene expression by the TYK2-related signal pathway, i.e., interferon α-dependent signaling. In summary, the activated TYK2/JAK1 heterodimeric complex could serve as a streamline factory as both an enzyme and a chaperone for converting the antiparallel uSTAT dimer to the parallel pSTAT dimers.

## Concluding Remarks

DEU is a TYK2-specific allosteric inhibitor that binds TYK2 JH2 but allosterically inhibits TYK2 JH1. Upon assembly and analysis of available structural information on TYK2, including the recently reported cryo-EM structures of activated (JAK1/IFNλR1)_2_ homodimeric complexes, we discovered that DEU and other TYK2 specific inhibitors exhibit a triple-action inhibitory mechanism. Specifically, they (i) stabilize the TYK2 JH1 autoinhibited state in complex with inhibitor-bound TYK2 JH1 through decreased protein dynamics of both domains, (ii) prevent binding of required ATP to TYK2 JH2 for the activation of TYK2 JH1, and (iii) destabilize the second active TYK2 JH1 complex with TYK2 JH1 for trans phosphorylation in the last step of activation via direct steric clashes with the bound inhibitor in TYK JH2. Additionally, we revealed a structural basis that could explain why TYK2-based interferon-induced gene expression is so rapid, and how the pSTAT1 homodimer and pSTAT1-pSTAT2 heterodimer are produced after activation during the JAK-STAT signaling pathway. The reported analysis of TYK2 has identified structural features that could facilitate new strategies for targeting the JAK-STAT signaling pathway with small molecule inhibitors.

## Materials and methods

Homolog modeling was carried out using both the CCP4 suite and the graphic program Coot based on secondary structure-based alignment (Emsley and Cowtan, 2004; Winn et al., 2011). Animation between two conformations was linearly extrapolated after transformation matrix was determined using the CCP4 suite (Winn et al., 2011). The extrapolation was made by dividing a total rotation around given axis into 20 or 30 steps alongside division of total skew translation along the axis. The direction of rotation was determined through visualization of linker topology and other geometric considerations as to whether it rotates by φ or by [2π-φ]. The starting and ending points of these two rotations are identical, but the rotation paths are different. Conversely, the usefulness of animation of this kind is to provide information as to which is the preferred path and sometimes is the only path of domain rotation during structural transition. We applied the simplest homolog modeling approach without any optimization in this study. Relevant coordinates used in this study are provided in Table 1.

## Funding

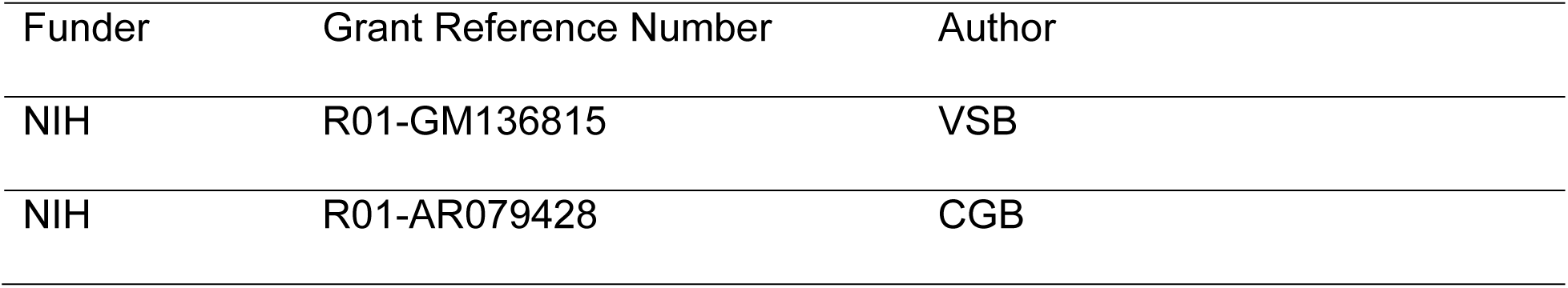

## Acknowledgements

This work was supported by the US National Institutes of Health (NIH)/National Institute of Arthritis and Musculoskeletal and Skin Diseases (NIAMS) under Award Number R01 AR079428 (to CGB) and by the US NIH under Award Number R01 GM136815 (to VSB). The content is solely the responsibility of the authors and does not necessarily represent the official views of the National Institutes of Health.

---

**Video 1.**
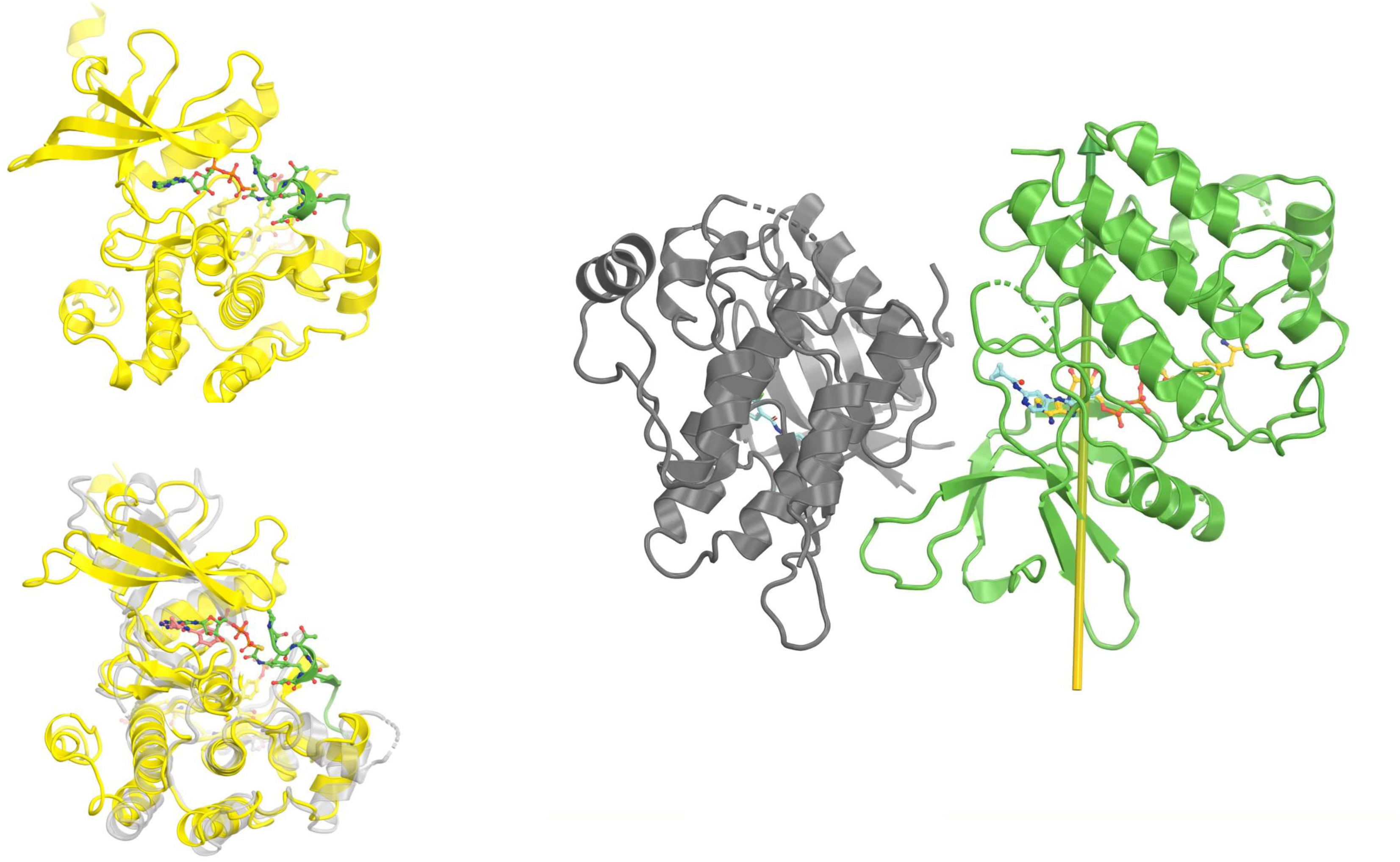

**Video 2.**
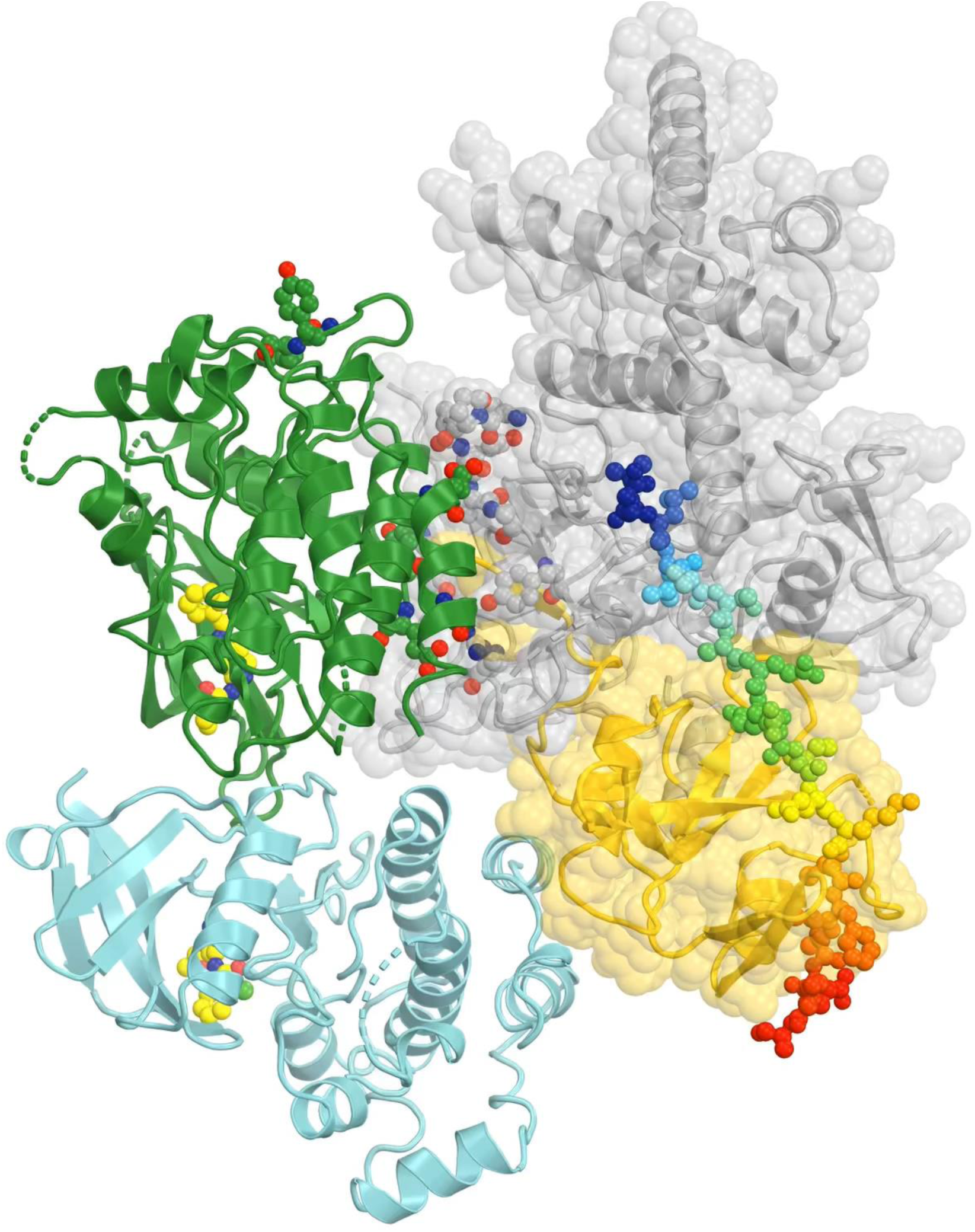

**Video 3.**
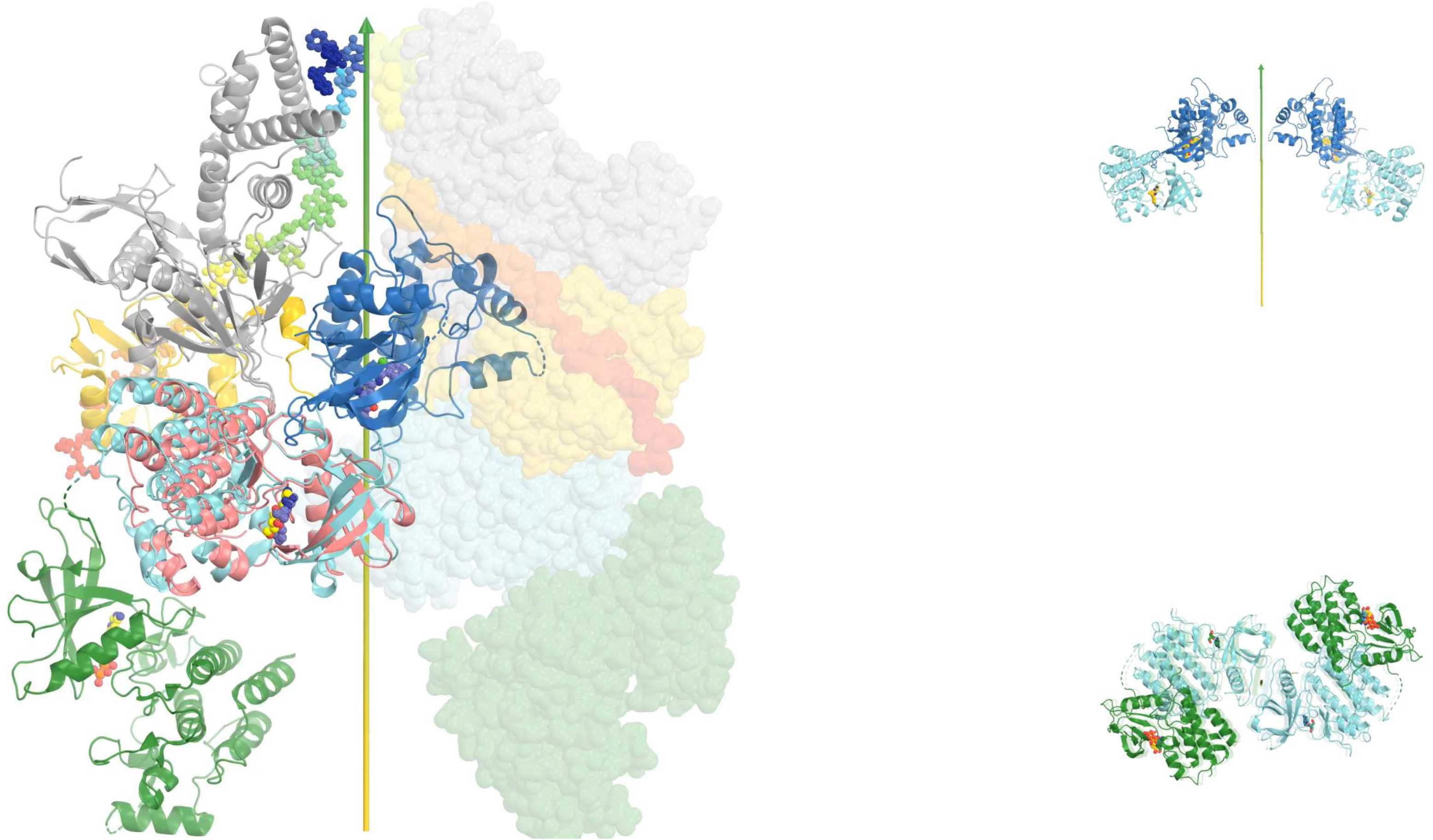

**Video 4.**
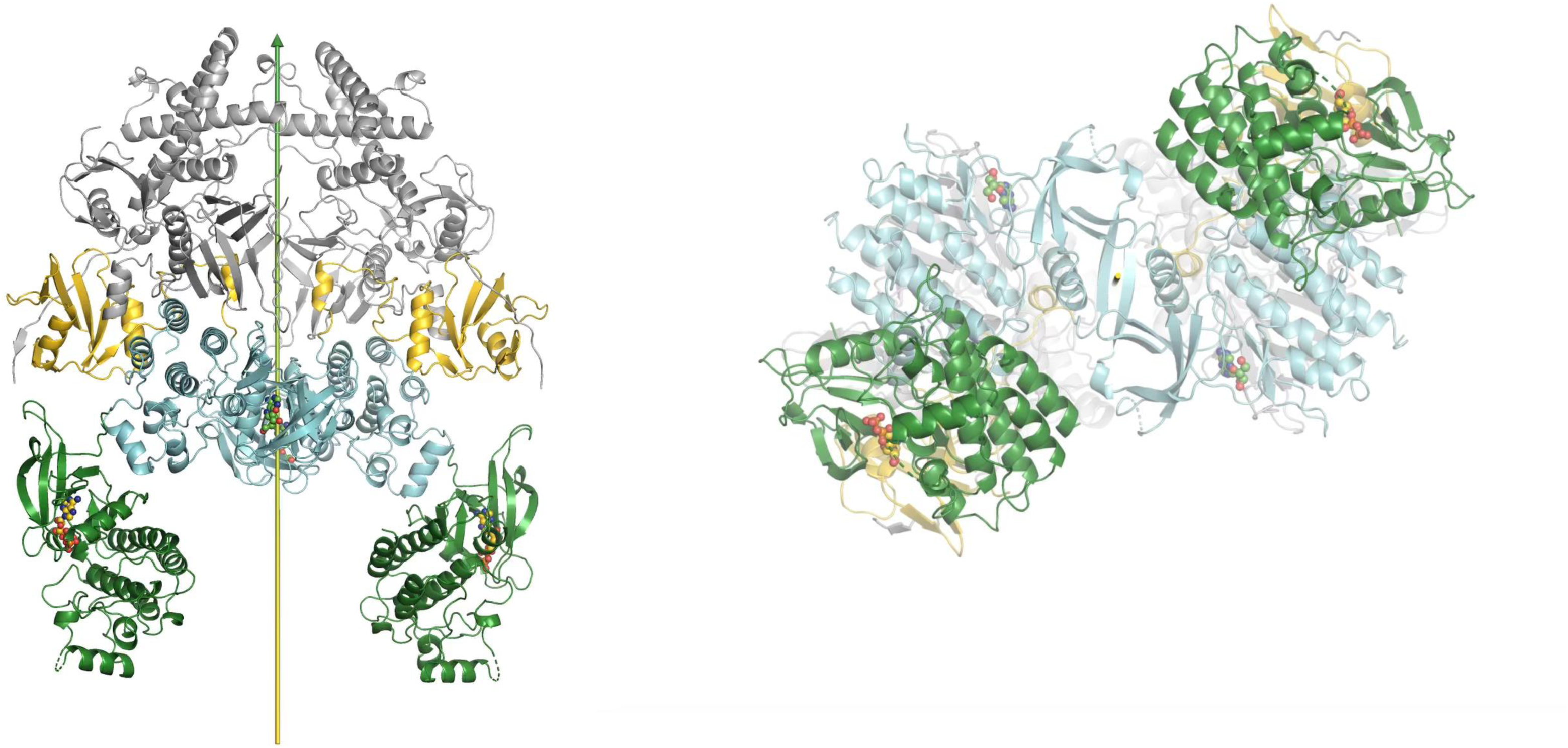

**Video 5.**
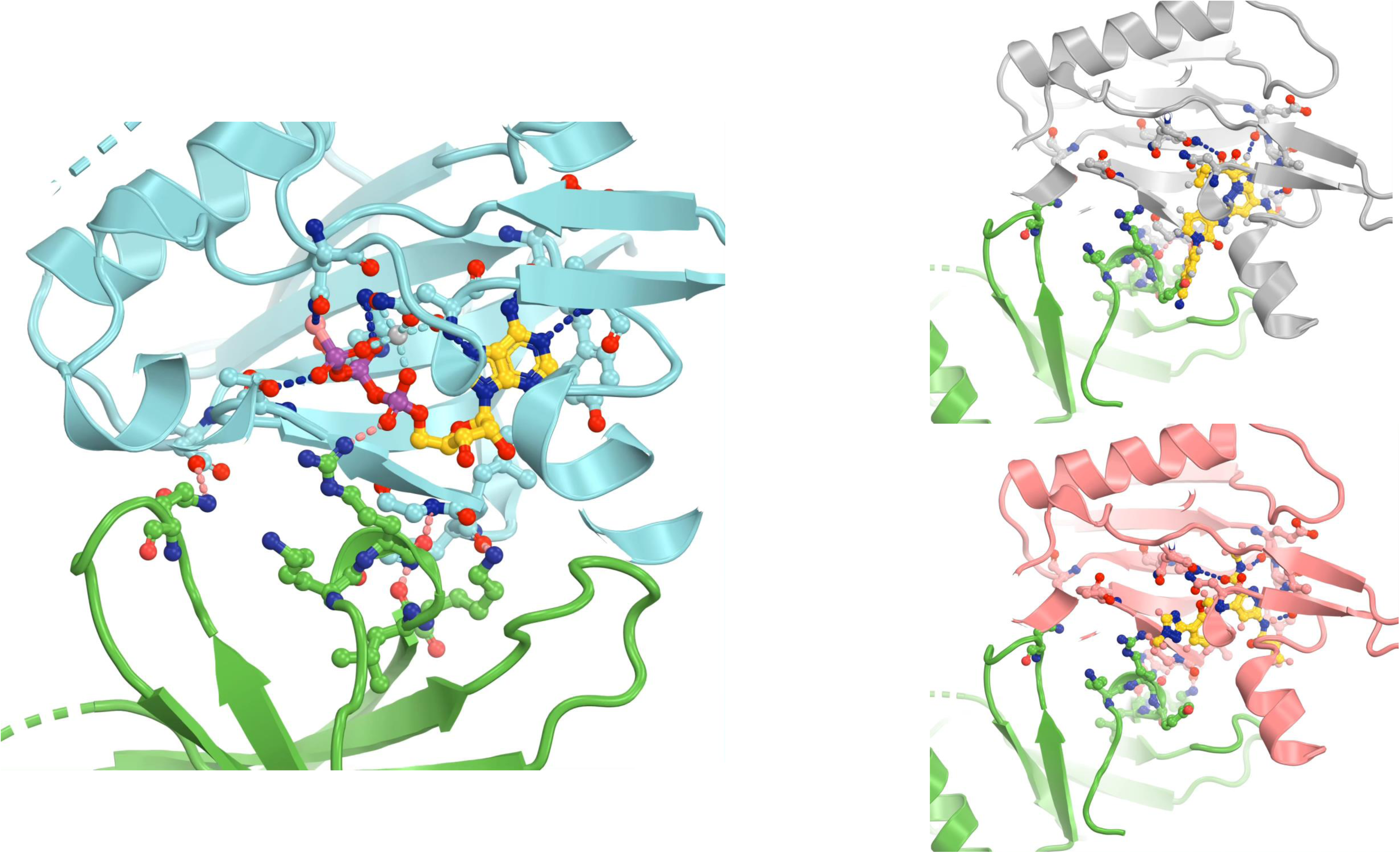

**Video 6.**
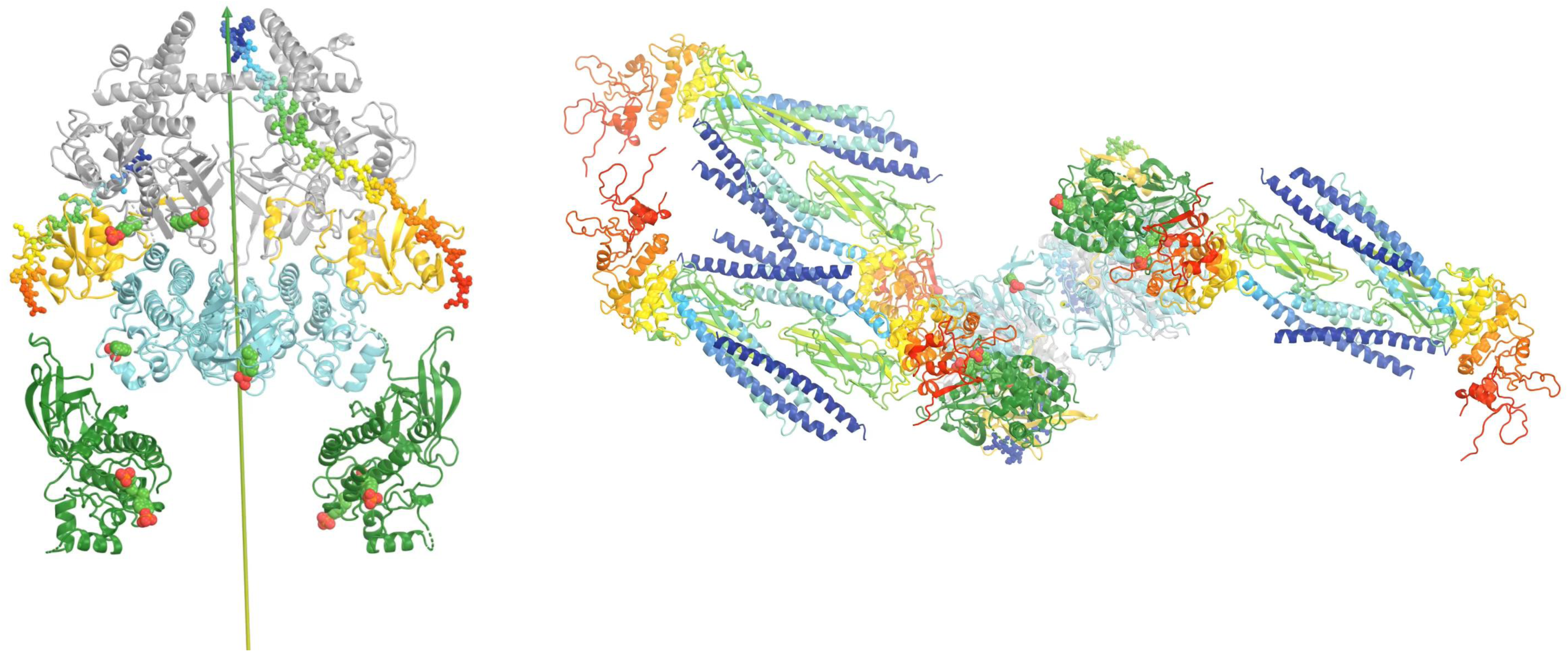

**Video 7.**
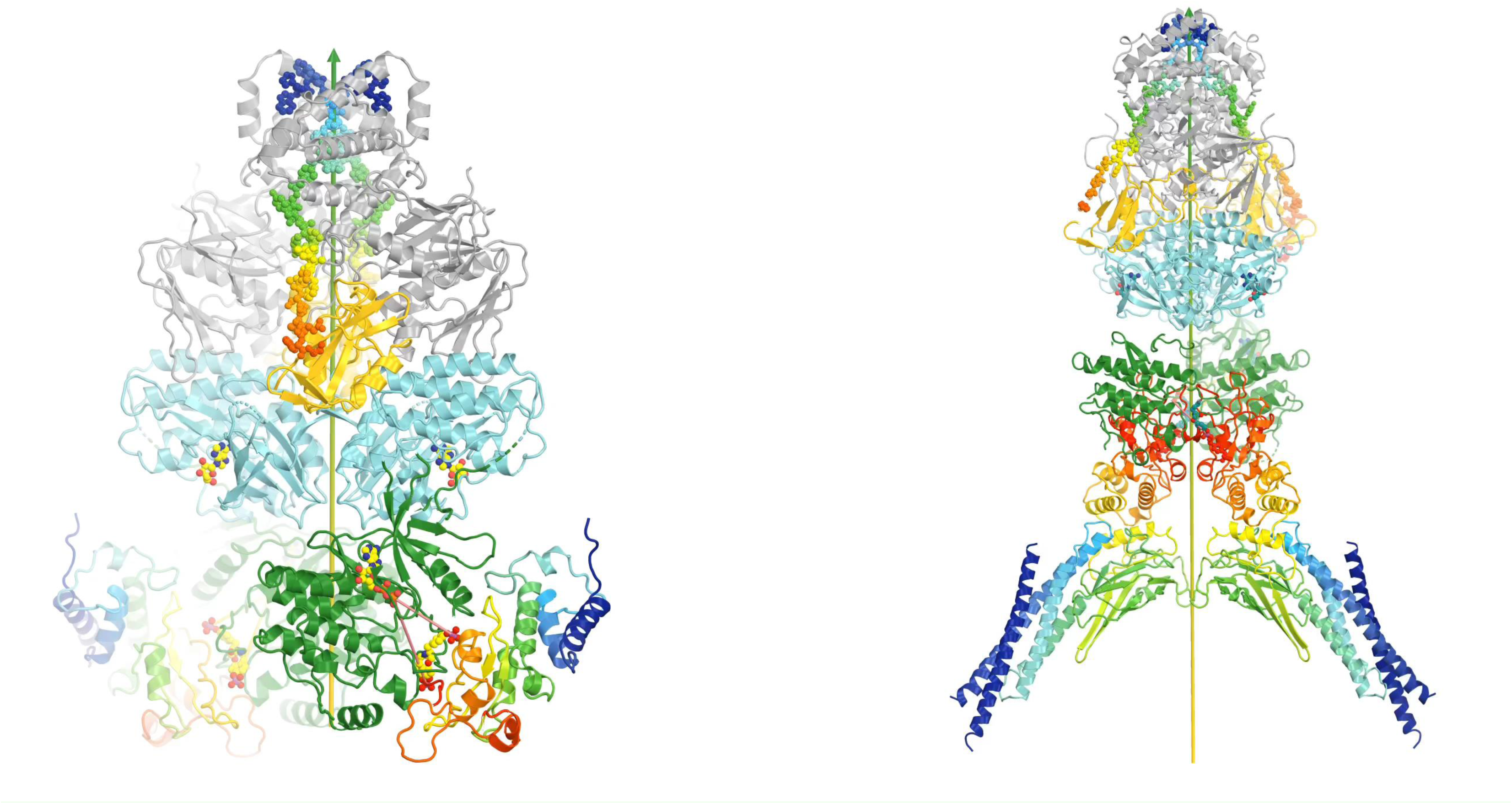

## References

Becker, S., Groner, B., and Muller, C.W. (1998). Three-dimensional structure of the Stat3beta homodimer bound to DNA. Nature 394, 145–151.

Burke, J.R., Cheng, L., Gillooly, K.M., Strnad, J., Zupa-Fernandez, A., Catlett, I.M., Zhang, Y., Heimrich, E.M., McIntyre, K.W., Cunningham, M.D., et al. (2019). Autoimmune pathways in mice and humans are blocked by pharmacological stabilization of the TYK2 pseudokinase domain. Sci Transl Med 11.

Caveney, N.A., Saxton, R.A., Waghray, D., Glassman, C.R., Tsutsumi, N., Hubbard, S.R., and Garcia, K.C. (2023). Structural basis of Janus kinase trans-activation. Cell Rep 42, 112201.

Chen, X., Vinkemeier, U., Zhao, Y., Jeruzalmi, D., Darnell, J.E., Jr., and Kuriyan, J. (1998). Crystal structure of a tyrosine phosphorylated STAT-1 dimer bound to DNA. Cell 93, 827–839.

Diallo, M., and Herrera, F. (2022). The role of understudied post-translational modifications for the behavior and function of Signal Transducer and Activator of Transcription 3. FEBS J 289, 6235–6255.

Emsley, P., and Cowtan, K. (2004). Coot: model-building tools for molecular graphics. Acta Crystallogr D Biol Crystallogr 60, 2126–2132.

Ernst, S., and Muller-Newen, G. (2019). Nucleocytoplasmic Shuttling of STATs. A Target for Intervention? Cancers (Basel) 11.

Firmbach-Kraft, I., Byers, M., Shows, T., Dalla-Favera, R., and Krolewski, J.J. (1990). tyk2, prototype of a novel class of non-receptor tyrosine kinase genes. Oncogene 5, 1329–1336.

Glassman, C.R., Tsutsumi, N., Saxton, R.A., Lupardus, P.J., Jude, K.M., and Garcia, K.C. (2022). Structure of a Janus kinase cytokine receptor complex reveals the basis for dimeric activation. Science 376, 163–169.

Greenlund, A.C., Farrar, M.A., Viviano, B.L., and Schreiber, R.D. (1994). Ligand-induced IFN gamma receptor tyrosine phosphorylation couples the receptor to its signal transduction system (p91). EMBO J 13, 1591–1600.

Hanks, S.K., Quinn, A.M., and Hunter, T. (1988). The protein kinase family: conserved features and deduced phylogeny of the catalytic domains. Science 241, 42–52.

Krolewski, J.J., Lee, R., Eddy, R., Shows, T.B., and Dalla-Favera, R. (1990). Identification and chromosomal mapping of new human tyrosine kinase genes. Oncogene 5, 277–282.

La Sala, G., Michiels, C., Kukenshoner, T., Brandstoetter, T., Maurer, B., Koide, A., Lau, K., Pojer, F., Koide, S., Sexl, V., et al. (2020). Selective inhibition of STAT3 signaling using monobodies targeting the coiled-coil and N-terminal domains. Nat Commun 11, 4115.

Li, J., Rodriguez, J.P., Niu, F., Pu, M., Wang, J., Hung, L.W., Shao, Q., Zhu, Y., Ding, W., Liu, Y., et al. (2016). Structural basis for DNA recognition by STAT6. Proc Natl Acad Sci U S A 113, 13015–13020.

Liu, C., Kieltyka, J., Fleischmann, R., Gadina, M., and O’Shea, J.J. (2021). A Decade of JAK Inhibitors: What Have We Learned and What May Be the Future? Arthritis Rheumatol 73, 2166–2178.

Liu, F., Wang, B., Liu, Y., Shi, W., Hu, Z., Chang, X., Tang, X., Zhang, Y., Xu, H., and He, Y. (2023). Design, synthesis and biological evaluation of novel N-(methyl-d(3)) pyridazine-3-carboxamide derivatives as TYK2 inhibitors. Bioorg Med Chem Lett 86, 129235.

Lupardus, P.J., Ultsch, M., Wallweber, H., Bir Kohli, P., Johnson, A.R., and Eigenbrot, C. (2014). Structure of the pseudokinase-kinase domains from protein kinase TYK2 reveals a mechanism for Janus kinase (JAK) autoinhibition. Proc Natl Acad Sci U S A 111, 8025–8030.

Manning, G., Whyte, D.B., Martinez, R., Hunter, T., and Sudarsanam, S. (2002). The protein kinase complement of the human genome. Science 298, 1912–1934.

Mao, X., Ren, Z., Parker, G.N., Sondermann, H., Pastorello, M.A., Wang, W., McMurray, J.S., Demeler, B., Darnell, J.E., Jr., and Chen, X. (2005). Structural bases of unphosphorylated STAT1 association and receptor binding. Mol Cell 17, 761–771.

Meyer, T., and Vinkemeier, U. (2004). Nucleocytoplasmic shuttling of STAT transcription factors. Eur J Biochem 271, 4606–4612.

Min, X., Ungureanu, D., Maxwell, S., Hammaren, H., Thibault, S., Hillert, E.K., Ayres, M., Greenfield, B., Eksterowicz, J., Gabel, C., et al. (2015). Structural and Functional Characterization of the JH2 Pseudokinase Domain of JAK Family Tyrosine Kinase 2 (TYK2). J Biol Chem 290, 27261–27270.

Neculai, D., Neculai, A.M., Verrier, S., Straub, K., Klumpp, K., Pfitzner, E., and Becker, S. (2005). Structure of the unphosphorylated STAT5a dimer. J Biol Chem 280, 40782–40787.

Parang, K., Till, J.H., Ablooglu, A.J., Kohanski, R.A., Hubbard, S.R., and Cole, P.A. (2001). Mechanism-based design of a protein kinase inhibitor. Nat Struct Biol 8, 37–41.

Philips, R.L., Wang, Y., Cheon, H., Kanno, Y., Gadina, M., Sartorelli, V., Horvath, C.M., Darnell, J.E., Jr., Stark, G.R., and O’Shea, J.J. (2022). The JAK-STAT pathway at 30: Much learned, much more to do. Cell 185, 3857–3876.

Spinelli, F.R., Meylan, F., O’Shea, J.J., and Gadina, M. (2021). JAK inhibitors: Ten years after. Eur J Immunol 51, 1615–1627.

Stark, G.R., and Darnell, J.E., Jr. (2012). The JAK-STAT pathway at twenty. Immunity 36, 503–514.

Tokarski, J.S., Zupa-Fernandez, A., Tredup, J.A., Pike, K., Chang, C., Xie, D., Cheng, L., Pedicord, D., Muckelbauer, J., Johnson, S.R., et al. (2015). Tyrosine Kinase 2-mediated Signal Transduction in T Lymphocytes Is Blocked by Pharmacological Stabilization of Its Pseudokinase Domain. J Biol Chem 290, 11061–11074.

Wallweber, H.J., Tam, C., Franke, Y., Starovasnik, M.A., and Lupardus, P.J. (2014). Structural basis of recognition of interferon-alpha receptor by tyrosine kinase 2. Nat Struct Mol Biol 21, 443–448.

Winn, M.D., Ballard, C.C., Cowtan, K.D., Dodson, E.J., Emsley, P., Evans, P.R., Keegan, R.M., Krissinel, E.B., Leslie, A.G., McCoy, A., et al. (2011). Overview of the CCP4 suite and current developments. Acta Crystallogr D Biol Crystallogr 67, 235–242.

Wrobleski, S.T., Moslin, R., Lin, S., Zhang, Y., Spergel, S., Kempson, J., Tokarski, J.S., Strnad, J., Zupa-Fernandez, A., Cheng, L., et al. (2019). Highly Selective Inhibition of Tyrosine Kinase 2 (TYK2) for the Treatment of Autoimmune Diseases: Discovery of the Allosteric Inhibitor BMS-986165. J Med Chem 62, 8973–8995.

Zhang, H., Cao, X., Tang, M., Zhong, G., Si, Y., Li, H., Zhu, F., Liao, Q., Li, L., Zhao, J., et al. (2021). A subcellular map of the human kinome. Elife 10.

